# Computational prediction of human deep intronic variation

**DOI:** 10.1101/2023.02.17.528928

**Authors:** Pedro Barbosa, Rosina Savisaar, Maria Carmo-Fonseca, Alcides Fonseca

## Abstract

The adoption of whole genome sequencing in genetic screens has facilitated the detection of genetic variation in the intronic regions of genes, far from annotated splice sites. However, selecting an appropriate computational tool to differentiate functionally relevant genetic variants from those with no effect is challenging, particularly for deep intronic regions where independent benchmarks are scarce.

In this study, we have provided an overview of the computational methods available and the extent to which they can be used to analyze deep intronic variation. We leveraged diverse datasets to extensively evaluate tool performance across different intronic regions, distinguishing between variants that are expected to disrupt splicing through different molecular mechanisms. Notably, we compared the performance of SpliceAI, a widely used sequence-based deep learning model, with that of more recent methods that extend its original implementation. We observed considerable differences in tool performance depending on the region considered, with variants generating cryptic splice sites being better predicted than those that affect splicing regulatory elements or the branchpoint region. Finally, we devised a novel quantitative assessment of tool interpretability and found that tools providing mechanistic explanations of their predictions are often correct with respect to the ground truth information, but the use of these tools results in decreased predictive power when compared to black box methods.

Our findings translate into practical recommendations for tool usage and provide a reference framework for applying prediction tools in deep intronic regions, enabling more informed decision-making by practitioners.

## Background

Genetic variation plays a crucial role in understanding human disease and trait inheritance Yet, for a long time, studies paid scant attention to variants in intronic gene regions [1], which were thought to harbor little functional variation. With the advent of whole genome sequencing (WGS), and the possibility to apply it at the population scale [2, 3], rare intronic variation can be identified at unprecedented levels. However, the sheer amount of candidate variants detected in the genome of an individual poses challenges for functional interpretation [4], particularly for variants affecting RNA splicing [5].

Splicing consists of removing introns from the primary transcript and is mediated by the spliceosome complex with the help of many RNA-binding proteins (RBPs) that recognize regulatory signals in exons and introns [6]. Splicing is tightly regulated across cell types and is sensitive to genetic variants occurring in *cis* (within the exons and introns of the splicing substrate) and in *trans* (within the genes encoding for splicing factors) [7]. It is estimated that 10 to 50% of all monogenic disease-causing variants affect pre-mRNA splicing [8, 9, 10]. In addition, cancer driver mutations are often associated with splicing alterations, notably in the case of trans variants that occur in genes encoding core components of the splicing machinery [11].

Of the cis variants that affect splicing, those that disrupt the canonical splice site elements (5ss, 3ss) have been studied the most thoroughly. Such variants are fairly easy to recognize because the splice site sequences are short and adhere to a highly conserved consensus [12]. Other “non-canonical” splicing variants can impact the binding of regulatory factors to splicing enhancers or silencers. These enhancers and silencers consist of short, poorly defined sequence motifs, which can occur at varying distances to the splice sites and can overlap either exons or introns [13]. It is thus difficult to identify them – and even more difficult to know when a mutation has disrupted them. Disruption of splicing information can lead to aberrant splice events such as exon skipping, full intron retention, or exon shortening or lengthening. Splice variants can also create entirely new exons (“pseudoexons”). This can happen when a mutation creates a novel splice site, as well as when an existing but inactive (“cryptic”) splice site is activated by the creation of an enhancer motif [14].

There have been continuous efforts to systematically catalog disease-causing variation in databases such as ClinVar [15] or the Human Gene Mutation Database (HGMD) [16]. These resources show enrichment of splicing-related variants in the vicinity of splice site regions. Partly, this reflects a biological reality, where the sequence around the splice sites is particularly dense in splicing-relevant information. However, this enrichment may in some measure also be due to the easier detection of splice site mutations, as well as a bias towards covering exons and splice sites in genetic screens [17]. Therefore, it is expected that many splicing variants in other gene regions remain to be discovered. Our dearth of knowledge is greatest deep inside the introns, where the detection problem is the hardest given the enormity of the search space and the fact that the rare splice-affecting variants are greatly outnumbered by mutations with no effect. As a result, deep intronic variants often end up labeled as Variant of Uncertain Significance (VUS) [18], although a subset may have great clinical importance. Indeed, recent evidence has shown that deep intronic mutations triggering pseudoexon activation are an overlooked cause of human disease [19, 20].

Given the additional challenges of interpreting deep intronic mutations, computational tools are often used to prioritize variants based on their likelihood of being deleterious. The first wave of methods used large genomics datasets to engineer features (e.g., allele frequencies from ExaC [21] or histone modification levels across cell lines from ENCODE [22]) and to build classifiers that work on tabular data. More recently, end-to-end deep-learning methods predict the impact of genetic variants from sequence alone, with the features automatically extracted within the network [23]. SpliceAI [10] is widely recognized as the most successful method of this kind, although its performance has been shown to vary across studies and datasets considered [5]. Recently, new models have been developed based on SpliceAI, either combining its predictions with other sources of information (such as genetic constraint for ConSpliceML [24] or tissue-specific splice site usage for AbSplice-DNA [25]) or creating an entirely new model based on SpliceAI architecture. For example, Pangolin [26] uses splicing quantifications from multiple species and tissues to not only predict whether a position is a splice site (as SpliceAI does) but also to predict splice site usage (e.g., how much a splice site is being used in a given tissue). In contrast, CI-SpliceAI [27] uses different training labels for true and false splice site positions based on a collapsed transcript structure derived from GENCODE [28] annotations.

Most intronic variant prediction benchmarking studies are performed by the authors of the tools to present a comparative analysis with existing methods. Even subconsciously, biases might be favoring the proposed model, be it because of the dataset selected or the methodology employed for the comparison [29, 30]. Multiple independent benchmark studies do exist [31, 32, 33, 34, 35, 36], however, their scope is often somewhat limited. Firstly, some studies only focus on variants overlapping particular types of splicing information, e.g. splicing regulatory elements [33]. Secondly, only using variants from a small number of genes can render the genome-wide extrapolation of conclusions difficult [32, 34]. Lastly, to our knowledge, no study compares the performance of promising and recently developed methods such as Pangolin, CI-SpliceAI, ConSpliceML, AbSplice-DNA, and SPiP [37].

To help researchers and clinical practitioners understand prediction tools and how they can be applied to interpret genetic variants in non-coding regions of genes, we conducted a comprehensive evaluation of a series of tools for the task of predicting functional variation in the intronic space far from canonical splice sites. To this end, we carefully selected intronic variants from multiple sources and curated a new set of disease-causing deep intronic variants affecting RNA splicing. Besides evaluating the capacity of tools to predict functional variants deep within the introns, we report, for the first time, an assessment of the interpretability of the output of these tools. We finally provide clear recommendations for tool usage depending on the variant’s location within the intron and its molecular effect.

## Results

### The prediction tools studied are diverse in methodology and objectives

In this study, we have provided a snapshot of the state-of-the-art of methods that predict, in any way, functional variation in introns (Table 1). We divided the methods into four different categories: conservation scores that measure the degree of evolutionary conservation at a given position or region of the genome; genome-wide predictors that integrate multiple feature types to predict variant effects regardless of the variant type; methods that focus on splice-disrupting variants and allow for automated batch predictions; and splicing-specific methods that solely target specific types of splicing information (e.g., Branchpoint (BP)), or require the use of a web application to retrieve results. For many of the tools, there are two fundamentally different ways to obtain predictions: making *de novo* model inferences given an input variant set or using pre-computed predictions, which is faster computationally. We decided to use pre-computed predictions when available because it considerably simplifies the variant annotation pipeline and is thus accessible to a more diverse set of end users. Out of the 37 tools used to score at least one dataset in this manuscript, 19 had pre-computed databases available (Table 1). Because some of them only provide predictions for the GRCh37 genome build, we ran all experiments using this genome version. Of note, pre-computed predictions are a permanent representation of a model version, which may not be updated along with developments to the tool. However, we observed that only one tool, CAPICE [38], had outdated pre-computed scores.

**Table 1:**
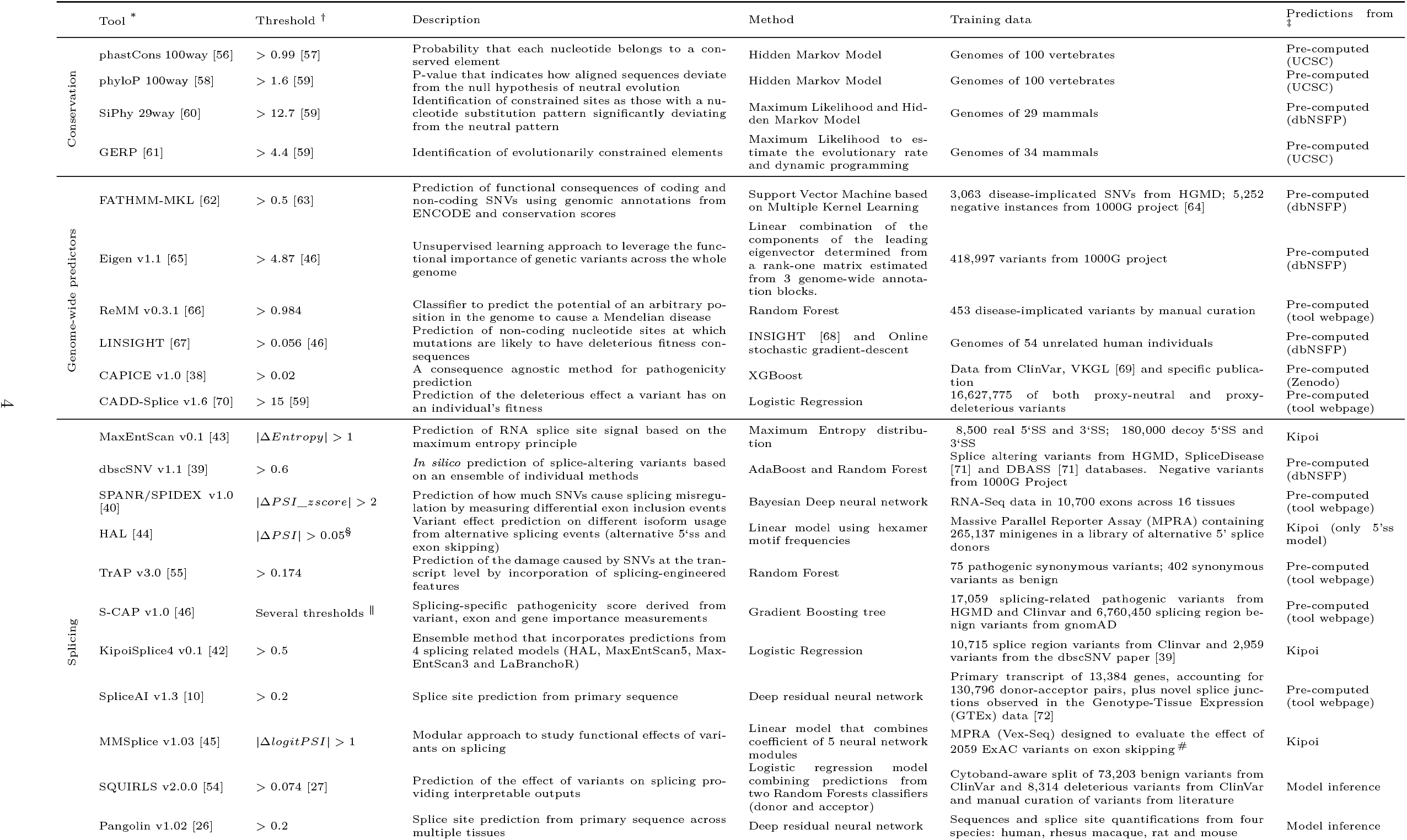
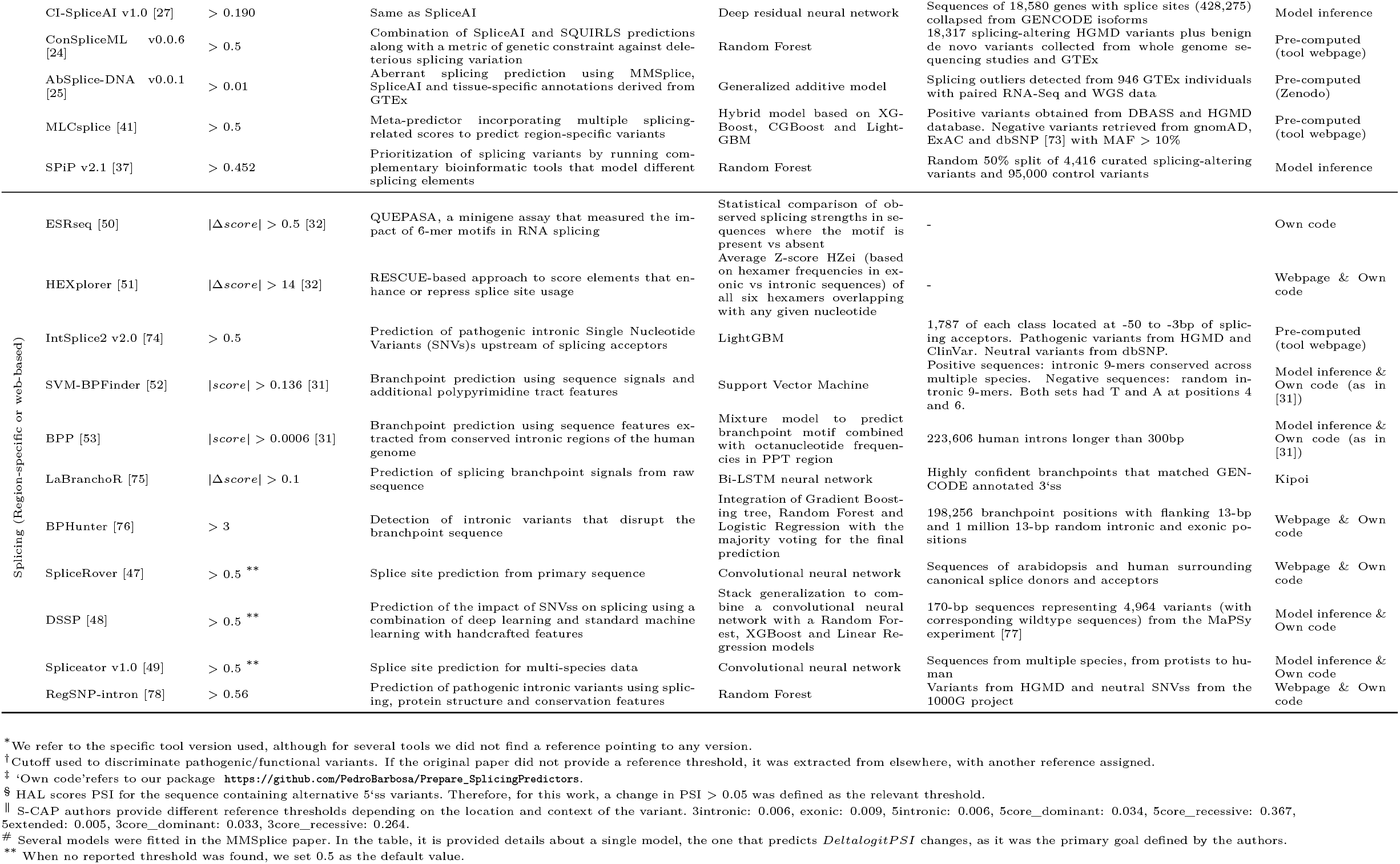
Summary of the computational methods used in this study

Importantly, not every tool considered was built with deep intronic regions in mind. For example, some were explicitly trained only to score near-splice site variants (e.g., dbscSNV [39]), while others only output predictions up to an approximately defined distance between the variant and the nearest splice site (e.g., 300 bp for SPIDEX [40] or 50 bp for MLCSplice [41]). In addition, we ran certain models (KipoiSplice4 [42], MaxEntScan [43], HAL [44], MMSplice [45]) using the Kipoi framework [42], which further restricts predictions to a tool-specific distance between the splice site and the variant. Therefore, we expected these methods to perform poorly on some comparisons simply because the fraction of missing predictions should increase when moving further into the intron. Still, we decided to include these tools in the study because many of the variants evaluated locate within the distance that we expected these tools to cover, especially for region-specific analyses.

It should also be noted that the tools were built for different tasks. While some models were designed to distinguish between pathogenic and benign variants (e.g., S-CAP [46], KipoiSplice4), others predict variant effects on splicing outcome, which does not necessarily translate into disease (e.g., SPiP, MMSplice). The latter category includes sequence-based deep learning models such as SpliceAI, which do not accept genetic variants as input. Rather, they predict the probability that a given sequence position functions as a splice site. By running the model with both the reference sequence and the mutated sequence, it’s possible to measure the effect of genetic variants on splice sites. If the model is run twice, once with the reference and once with the mutated sequence, it is possible to measure splice site alterations caused by genetic variants. This has been the major practical use of the tool so far. Using the same approach, we also included several sequence-based methods that predict splicing-related elements. These include SpliceRover [47], DSSP [48], and Spliceator [49] for splice site-associated variants, ESRseq [50] and HEXplorer [51] for variants affecting splicing regulatory elements, and SVM-BPFinder [52] and BPP [53] for variants impacting the BP signal. Of note, we only employed these methods for the datasets deemed to be relevant given their original task.

### Intronic pathogenic variants located beyond 10 bp from the splice sites are poorly predicted

We employed a bin-based analysis to evaluate ClinVar data (Supplementary Table S1). Because ClinVar contains disease-causing variants that act through different molecular mechanisms, we included not only splicing-related tools but also conservation scores and whole genome predictors in the evaluations. As expected, the distribution of the two variant classes (pathogenic and benign) across bins is highly unbalanced (Figure 1A). More than 90% of the intronic pathogenic variants occur at splice site positions, and more than 95% occur within 10 nucleotides from an exon-intron boundary.

**Figure 1:**
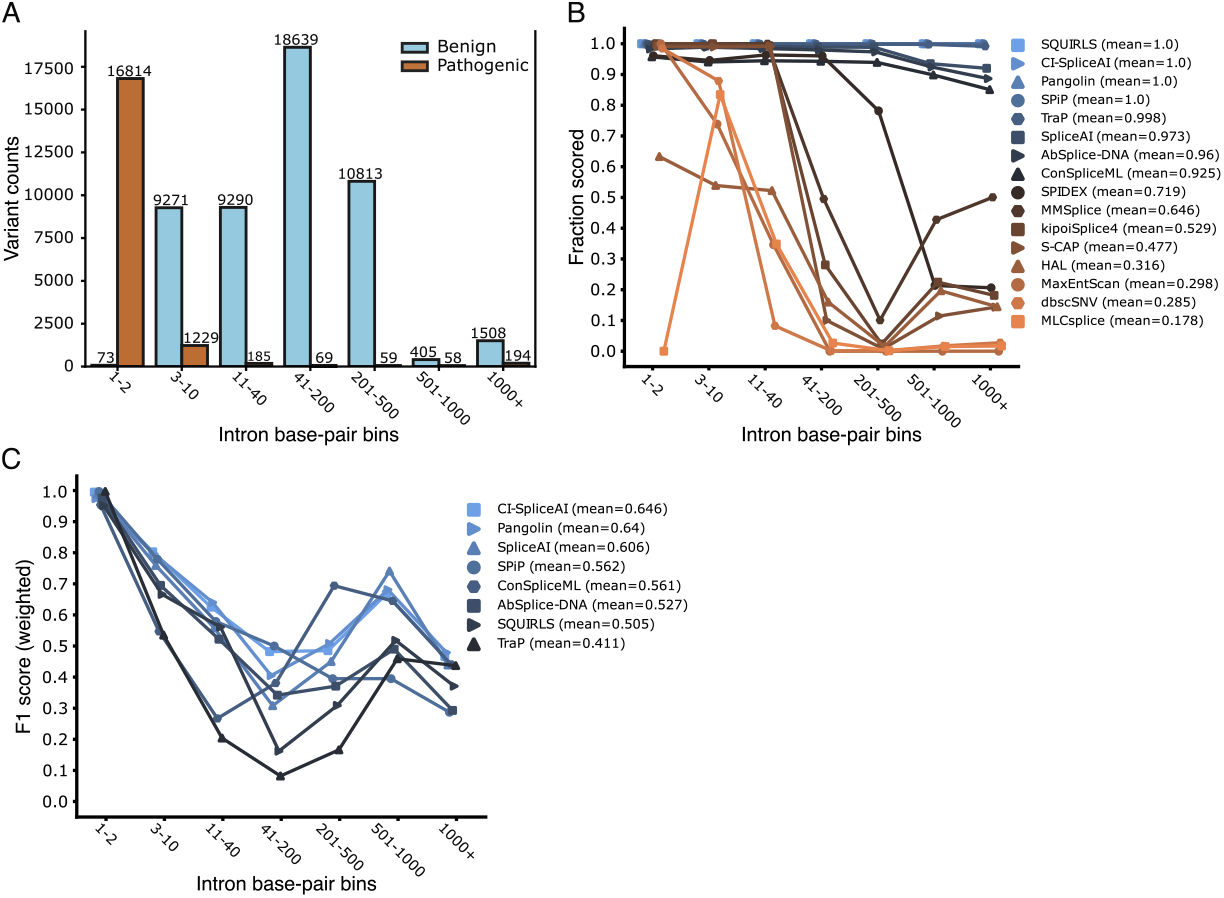
Intronic variant prediction in ClinVar. **A** - Distribution of ClinVar variants across each intronic bin. **B** - Fraction of variants scored (predicted) at each intronic bin for splicing-associated methods. Mean values in the legend represent the average fraction of variants scored across all bins. **C** - Performance of splicing-associated tools that predict entire introns at each intronic bin. Mean values in the legend represent the average weighted F1 score across all bins.

Because of the spatial limitations discussed previously, we expected that some tools would only output predictions for ClinVar variants located close to splice sites. Our results confirmed that several methods make predictions for less than 50% of the variants located at a distance of more than 40 bp from the nearest splice junction (Figure 1B). The fraction of predicted variants decreases according to the expected regions that each model covers: 50bp for S-CAP and MLCsplice, and 300bp for SPIDEX. MLCsplice was designed to predict non-canonical splicing variants (i.e. excluding splice site variants), thus, it is the only tool that displays no predictions at 1-2 positions (Figure 1B). In addition, we observed that the tools run using the Kipoi framework (KipoiSplice4, MaxEntScan, HAL, MMSplice) displayed a notable drop 41-200 bp from the splice site. On the other hand, SQUIRLS [54], Pangolin, CI-SpliceAI, SPiP, TraP [55], SpliceAI, ConSpliceML and AbSplice-DNA predict across entire introns (Figure 1B). Regarding the remaining tool categories, we observed that both whole genome predictors and conservation scores (except phastCons [56]) output predictions for most ClinVar variants (Supplementary Figure S1A, C).

Next, we evaluated how the tools that score across full introns perform with ClinVar data. Performance dropped considerably for variants located deeper in intronic regions, especially once a distance of 10 nucleotides from the splice site had been reached (Figure 1C, Supplementary Figure S1B, D). The splicing tools with the smallest and largest performance drop between the splice site bin (“1-2”) and the “11-40” bin were Pangolin and TraP, with weighted F1 scores decreasing by 0.334 and 0.793, respectively (Supplementary Table S2, Figure 1C). Conservation scores and whole-genome predictors performed poorly as well. Except for CAPICE and CADD-Splice, most methods displayed weighted F1 scores below 0.1 at the 11-40 bin (Supplementary Figure S1B, D). Overall, the most performant tools were CAPICE, CI-SpliceAI and Pangolin with an average weighted F1 score across all intronic bins of 0.70, 0.65 and 0.64, respectively (Figure 1C, Supplementary Figure S1B). Because CAPICE included ClinVar data in model training, we addressed circularity type I problems [15] by removing the ClinVar variants used to train CAPICE. 14,491 variants were excluded (5,259 pathogenic and 9,232 benign), resulting in 54,117 variants to evaluate. As a result, CAPICE performance dropped 10 units to an average weighted F1 score of 0.60 (Supplementary Figure S1E). In addition, we inspected performance using Precision-Recall Curves (PR Curves), since the weighted F1 metric measures performance based on a single pre-defined threshold to separate the pathogenic from the benign classes, which may not be tuned to predict clinically relevant intronic variants. Indeed, performance values improved for some tools, suggesting that the reference thresholds used are suboptimal (Supplementary Figure S1 F).

Strikingly, we noticed an increase in performance in the deepest intronic bins when compared to intermediate distances (Figure 1C, Supplementary Figure S1B, D). We hypothesized that variability in transcript structures could be the reason: despite these variants being assigned as occurring very deep within introns (> 500bp from the splice site of the canonical isoform) in the reference isoform, they may be exonic or near-splice site variants of other isoforms of the associated gene. To tackle this question, we looked at the raw transcript annotations (by running Ensembl Variant Effect Predictor (VEP) without picking consequences) of the variants assigned to the > 500bp bin (N=2165) and decomposed them into several sub-categories based on their localization in different transcript isoforms (see Methods). Our analysis revealed that 742 variants are located in exons in other transcripts and 299 variants mapped to introns but closer to splice sites than in the transcript isoform originally considered (Supplementary Figure S2A). In particular, some of the intronic variants are located at splice sites in other transcripts (Supplementary Figure S2B). We found that the performance of the tools was generally better for these categories than for categories where the variant distance to the splice site remained unchanged (Supplementary Figure S2C), which is consistent with the hypothesis that deep intronic pathogenic variants are hard to predict. After excluding variants from exonic and closer-to-splice sites categories, we repeated the per-bin analysis to see whether the performance increase in the deepest bins remained. We observed that conservation-based methods and whole genome predictors displayed a decline in performance compared to the original analysis. On the other hand, a subset of splicing tools such as Pangolin or CI-SpliceAI showed better performance than before (Supplementary Figure S2D), suggesting that unequivocal deep intronic variants in ClinVar are associated with splicing and that SpliceAI-based methods can capture them reasonably well.

### Pathogenic splicing-affecting variants are captured well by deep learning based methods

Not all ClinVar intronic variants are associated with splicing defects. However, splicing-related tools were the most successful at predicting the pathogenicity of deep intronic mutations. Therefore, we decided to narrow our focus to variants that specifically affected splicing. We previously published a dataset of deep intronic variants causing human disease via disruption of splicing [19]. In the current study, we extended this dataset by performing a comprehensive literature search for case reports published after 2017, where the association between a variant and a splicing defect was supported by experimental evidence, such as from RT-PCR, sequencing of cDNA products, RNA-Seq or minigene/midigene assays (Supplementary Table S3). The dataset is composed of 162 variants covering a diverse range of disease phenotypes, with most diseases represented by fewer than 3 variants (Supplementary Figure S3A). A great number of these variants are not yet reported in ClinVar (N=88), and of those that are reported, a few (N=13) are incorrectly classified as benign or as VUS (Supplementary Figure S3B). As further evidence of their pathogenicity, the variants are very rare in the general population, as most of them are absent from gnomAD [2], a widely used catalog of genetic variation across human populations (Supplementary Figure S3B, C).

The results showed that SpliceAI-derived methods outperformed the remaining tools. ConSpliceML displayed the highest area under the ROC (auROC), followed by SpliceAI, Pangolin and CI-SpliceAI (Figure 2A). However, evaluation using single thresholds revealed lower performance than using auROC, which is based on multiple thresholds (Supplementary Figure S3D). As practical clinical applications usually require a binary decision, this prompted us to optimize reference thresholds for detecting splice-affecting pathogenic intronic variation outside of the canonical splice site regions (see Methods). After threshold recalibration, we reveal SpliceAI and Pangolin as the best tools (weighted normalized MCC > 0.90) to identify pathogenic variants using a single cutoff value (Figure 2B, Supplementary Table S4). As a practical outcome of this analysis, we provide recalibrated thresholds for different trade-offs between precision and recall (Supplementary Table S5). When available, we recorded information on the molecular consequences of each variant on splicing. Pseudoexon activation was the most frequent consequence of deep intronic variants (193 out of 243 in our dataset). We also identified 39 variants leading to partial intron retention due to the usage of an alternative splice site. Exon skipping was observed in only 6 cases, consistent with previous observations that functional deep intronic variants are less commonly linked to this mechanism [79]. We next compared the tools’ ability to detect pseudoexon activation and partial intron retention variants using the optimized thresholds. We hypothesized that the tools would perform better on the latter group since these variants are located closer to the splice sites than those that activate pseudoexons (Supplementary Figure S3E). Nonetheless, we observed no statistically significant differences between the two groups (Figure 2C).

**Figure 2:**
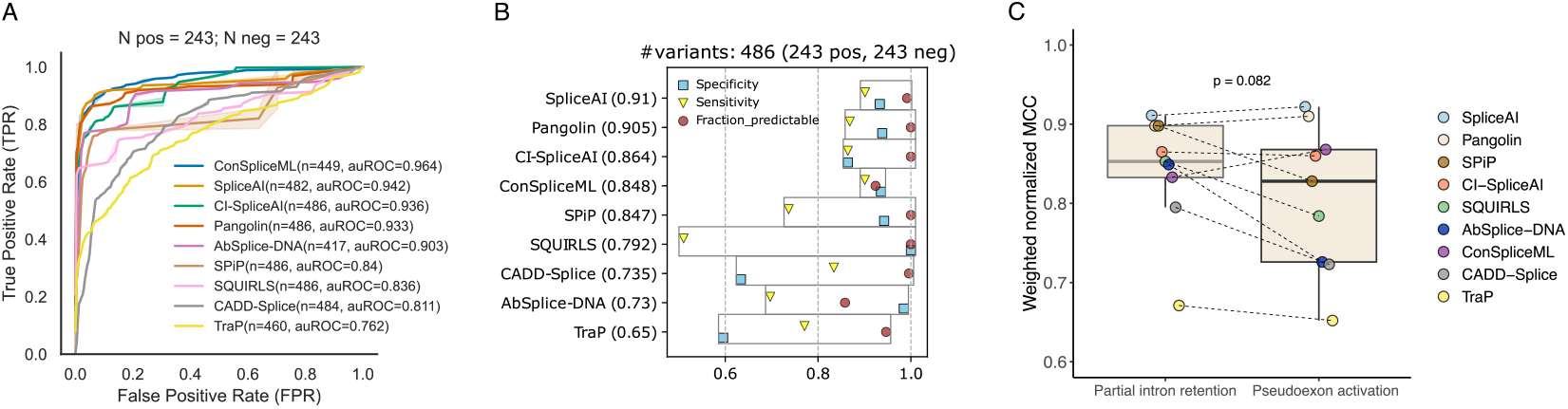
Pathogenic variant prediction of deep intronic variants affecting RNA splicing (81 variants from [19] and 162 curated for this manuscript). **A** - Receiving Operating Characteristic (ROC) analysis for all splicing-associated methods. **B** - Performance using optimized thresholds for intronic variation outside of canonical splice regions. The weighted normalized MCC was used to rank the tools. **C** - Performance using optimized thresholds on two different subsets of variants: variants leading to partial intron retention and variants leading to pseudoexon activation. Wilcoxon Signed Rank test, performed as a one-sided test, was used to compare the values between the two groups of variants.

### Performance varies considerably when predicting splicing-altering variants associated with different molecular mechanisms

To gain further insight into the molecular mechanisms driving the splicing alterations, we generated datasets of alternative splicing events triggered by intronic variants occurring at different regions that are important for splicing regulation (Table 2). We defined six categories (Figure 3A, see Methods). Within each region, we separately evaluated variants that trigger partial intron retention (via alternative splice site usage at annotated exons) and variants that lead to pseudoexon activation. Importantly, contrary to the analyses performed above, we evaluate performance not based on the ability to distinguish between pathogenic and non-pathogenic variants but rather between variants that do (positive class) or do not (negative class) affect the mechanistic splicing outcome. This is relevant as a variant may create a pseudoexon, thus affecting the outcome of splicing, without necessarily leading to disease. This decision was made as information on variant pathogenicity was not always available. We also switch to reporting performance using the Average Precision (AP) score as some categories have unbalanced data. Moreover, this scoring method, unlike F1 and MCC-based metrics, does not require us to define a single cut-off that could be applied to all categories of variants, which may have different optimal thresholds.

**Table 2:** Sources of data used to build region-specific splicing datasets.

| Study | Variants* | Per category** | Description |
| --- | --- | --- | --- |
| *Splicing-altering* |  |  |  |
| Vaz-Drago et al. [19] | 81 | AU=3;NSA=12;EL=9;<br>NSD=42;DD=15 | Manual curation of disease-causing deep intronic variants with experimental validations |
| Our curation | 141 | BP=9;AU=12;NSA=26;<br>EL=17;NSD=45;DD=32 | Manual curation of disease-causing deep intronic variants with experimental validations |
| Keegan et al. [20] | 153 | BP=3;AU=12;NSA=20;<br>EL=15; NSD=73;DD=30 | Characterization of hundreds of mutation events driving cryptic splicing via pseudoexon activation |
| Petersen et al. [80] | 12 | BP=1;AU=1;EL=9<br>;DD=1 | Characterization of pseudoexons activated by deep intronic mutations that do not create or strengthen cryptic splice sites |
| Tubeuf et al. [32] | 3 | EL=3 | Benchmark of user-friendly tools for predicting variants affecting splicing regulatory elements |
| Jung et al. [79] | 252 | BP=48;AU=40;NSA=4;<br>EL=51;NSD=41;DD=68 | Identification of intronic mis-splicing mutations from RNA-Seq using a read ratios approach |
| Moles-Fernández et al. [33] | 17 | BP=1;NSA=3;EL=2;<br>NSD=9;DD=2 | Benchmark of using both SpliceAI and user-friendly tools to identify deep intronic variants that disrupt splicing |
| *Splicing-neutral* |  |  |  |
| Moles-Fernández et al. [33] | 99 | BP=2;AU=36;DD=61 | Benchmark of using both SpliceAI and user-friendly tools to identify deep intronic variants that disrupt splicing |
| Vex-seq [81] | 283*** | BP=59;AU=51,<br>EL=123;DD=50 | Vex-seq, a MPRA to test the impact of 2059 variants in splicing across 110 alternative exons |
| MFASS [82] | 109 | BP=34; AU=17; DD=58 | Multiplexed functional assay (MFASS) that assayed the splicing effect of 27,733 ExAC variants |
| gnomAD [2] | 278 | NSA=67; NSD=211 | Common (and hypothetically benign) variants that create true splice site motifs |

\* Total number of variants used from original study. Since several variants were duplicated across studies, we kept unique occurrences given the order they appear in the table (top to bottom).
\*\* Number of variants contributing to each category. BP = Branchpoint-associated; NSA = New splice acceptor; NSD = New splice donor; AU = Acceptor upstream; DD = Donor downstream; EL = Exonic-like. Note: we could not assign a category to 21 variants of our curation as well as 2 variants from Moles-Fernández et al. [33], thus these variants were excluded.
\*\*\* Exceptionally, 123 variants from this study are exonic.

**Figure 3:**
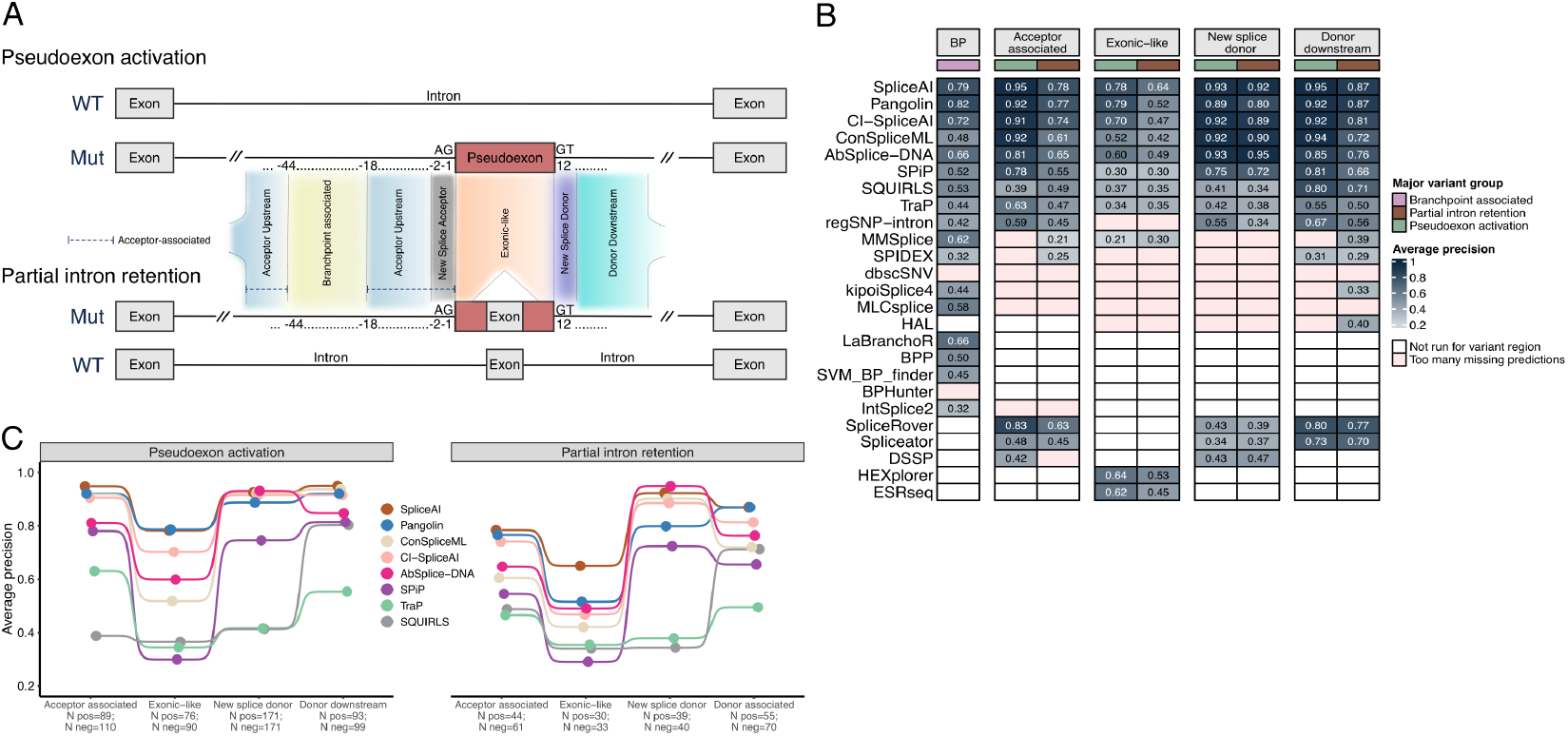
Tool performance evaluation on multiple regions associated with the regulation of splicing. **A** - Schematic representation of the regions used to define each dataset. For each of the two major groups (pseudoexon activation and partial intron retention), we show the expected wildtype (WT) structure in the absence of the variant, as well as the abnormal structure caused by the variant (Mut). Red blocks represent regions of the mRNA that are incorrectly spliced in. Exceptionally, some branchpoint-associated variants result in exon skipping, which is not graphically represented in the figure. **B** - Global overview of the performance for all the datasets analyzed. **C** - Average precision scores for the tools that can predict entire introns.

### Branchpoint-associated variants

Branchpoint-associated variants were defined as located 18 to 44 bp upstream of the splice acceptor of a cryptic or canonical splice site, either leading to pseudoexon activation, partial/full intron retention, or exon skipping. We collected most of the positive branchpoint variants from Jung et al. [79] (Table 2). The negative variants were located 18 to 44 bp upstream of an annotated splice site and had been shown not to affect splicing using the minigene-based reporter assays Vex-seq [81] and MFASS [82](Table 2, Supplementary Table S6). Because variants leading to pseudoexon inclusion were scarce (N=7) and the molecular consequences are not clear-cut (e.g., the same BP variant may lead to intron retention and to exon skipping), we analyzed all the variants affecting the BP region together. For this analysis, we additionally included four branchpoint prediction tools: SVM-BPFinder, BPP, LaBranchoR [75] and BPHunter [76]. Moreover, we included IntSplice2 [74] and regSNP-intron [78] since they predict splicing-associated SNVs at intronic positions overlapping the branchpoint region.

Deep learning-based methods displayed the highest AP for branchpoint-associated variants, with Pangolin, SpliceAI and CI-SpliceAI achieving 0.817, 0.786 and 0.716 respectively (Figure 3B, Supplementary Figure S4A). The best branchpoint-specific tool was LaBranchoR, with an AP of 0.661, followed by BPP (AP of 0.503, Supplementary Table S7). Unfortunately, BPHunter only reported the variants predicted to disrupt the BP, rendering the Precision-Recall Curves (PR Curves) analysis impossible. We further investigated whether there was any added value to using any tool besides Pangolin or SpliceAI. For the 22 variants (17 splicing-altering, 5 non-splicing-altering) that neither Pangolin nor SpliceAI predicted correctly using the adjusted thresholds, TraP, ConSpliceML, LaBranchoR and BPP displayed the best sensitivity (Supplementary Figure S5A). However, these 4 methods are likely to increase the false positive rate, as for the full dataset, specificity was lower than sensitivity (Supplementary Table S7). When we trained a Random Forest model on the full BP dataset using the predictions of the tools as features, we observed that LaBranchoR was an important feature of the model (Supplementary Figure S5B). Despite the limitations of having such a limited amount of data to train on, our results revealed LaBranchoR as the single branchpoint-specific tool to combine with the most performant generic tools, such as Pangolin or SpliceAI.

### Acceptor-upstream and new splice acceptor variants

The “acceptor-upstream” category refers to splicing-altering variants that locate upstream (up to 18bp) of an existing cryptic splice acceptor and activate it. On the other hand, the “new splice acceptor” category contains variants that form new splice sites themselves (Figure 3A). We collected negative variants differently for each of the two categories. For “acceptor-upstream” variants, we extracted variants located upstream of annotated splicing acceptors that did not interfere with the splicing outcome, as demonstrated through MFASS, Vex-seq or Moles-Fernández et al. [33]). The BP region from 18 to 44 bp was excluded. Conversely, we assigned common deep intronic gnomAD variants that create new splice acceptor motifs as negative “new splice acceptor” variants (Table 2). Because of their high frequency in the general population, it is assumed that these variants, although creating a splice acceptor motif, do not create a functional splice acceptor (see Methods).

The sets of splicing-altering variants we collected for each category were similar in size (70 and 67 variants for acceptor-upstream and new splice acceptor, respectively). However, when we split the variants according to the major molecular group (pseudoexon inclusion vs. partial intron retention), we obtained a very small number of new splice acceptor variants in the partial intron retention group (N=13), hence rendering their computational evaluation difficult. Therefore, for this particular analysis, we merged acceptor-upstream and new splice acceptor variants into a new “acceptor-associated” class so that we could have a reasonably large dataset to evaluate (Supplementary Table S6). As for the branchpoint-associated variants, we added IntSplice2 and regSNP-intron to the list of tools to evaluate. In addition, we included three splice site prediction methods that we customized to predict variant effects in VCF format: SpliceRover, DSSP and Spliceator.

SpliceAI, Pangolin, ConSpliceML and CI-SpliceAI achieved good performance on pseudoexon-activating variants, with AP above 0.9 (Figure 3B). However, for variants causing partial intron retention, performance dropped considerably, with no tool producing AP scores larger than 0.8 (Figure 3B). Nevertheless, the top tools remained the same, with SpliceAI, Pangolin and CI-SpliceAI displaying AP scores of 0.784, 0.766 and 0.741, respectively (Supplementary Figure S4C, Supplementary Table S7). Among the tools specifically added for this analysis, SpliceRover was the most competitive, ranking fifth for both pseudoexon and partial intron retention variants (Supplementary Figure S4B, C).

### Exonic-like variants

We consider here intronic variants that lie within either an activated pseudoexon or within an annotated exon that undergoes alternative splice site usage (Figure 3A). We identified 107 splicing-altering variants to compare against 123 splicing-neutral exonic variants from Vex-seq (Supplementary Table S6). After grouping the variants according to the major group, we obtained 76 pseudoexon-activating variants vs. 30 variants triggering partial intron retention. Accordingly, we randomly split negative the variants between the two groups so that the final datasets were fairly balanced (90 and 33 variants for each group, respectively). For this comparison, we also included two approaches that quantify splicing regulatory elements that enhance or repress flanking splice sites: ESREseq scores and HEXplorer.

Once again, we observed better overall performance for the pseudoexon group compared to the partial intron retention group (Figure 3B, Supplementary Figure S4D, E). Pangolin and SpliceAI were among the best tools in both major groups, yet SpliceAI displayed the best performance for the partial intron retention dataset by a large margin (> 0.12 AP difference to the second-best-ranked tool). Interestingly, HEXplorer and ESREseq performed better than several models that incorporate deep learning based predictions such as MMSplice, AbSplice-DNA or ConSpliceM. HEXplorer even displayed better performance than Pangolin (a pure sequence-based model that uses a much larger sequence context) for the partial intron retention dataset (Supplementary Figure S4E, Supplementary Table S7).

Although SpliceAI performed reasonably well, its pre-computed scores were configured to only report variant effects in a 50-bp window from the variant site. While this window is fine for most variant types (the affected splice sites are usually close to the variant site), that may not be the case for pseudoexon-activating variants that could be located deep inside the pseudoexon (assuming a pseudoexon of the size of an annotated exon). Therefore, we selected the splicing-altering variants missed by SpliceAI using the optimized threshold (N=30) and used the SpliceAI Lookup API(https://spliceailookup.broadinstitute.org/; last accessed November 30th, 2022) to run the model using a larger maximum distance (200 bp). We observed that 9 out of 30 were correctly reclassified as splicing-altering (Supplementary Table S8), suggesting that SpliceAI performance may be underestimated when ignoring longer-range variant effects.

### New splice donor variants

We identified 211 positive variants falling into this category (Table 2). For the negative set, we used variants that created a GT dinucleotide resulting in a splice donor consensus (GGTAAG), but that were unlikely to act as a cryptic splice site as they appeared in gnomAD with a population frequency >5% and were not observed to be used as a splice junction in GTEx [72] individuals. We again added SpliceRover, DSSP, Spliceator and regSNP-intron tools to the evaluation.

AbSplice-DNA performed best for both pseudoexon-activating and partial intron retention variants, with AP scores of 0.931 and 0.949, respectively (Supplementary Figure S4F, G), followed by SpliceAI and ConSpliceML, all of them with a performance metric above 0.9 (Figure 3B, Supplementary Table S7). Importantly, we noticed a large performance gap between SpliceAI-related tools (plus SPiP) and the rest, which performed rather poorly (almost all tools with AP scores below 0.5, Figure 3B, Supplementary Figure S4F, G). Considering that splicing-negative variants in this dataset create hypothetical splice donor decoys, we wondered whether tools that incorporate cryptic splice site scoring features using short sequence windows surrounding the variant site (Position Specific Scoring Matrix (PSSM)-based for regSNP-intron and TraP, information content-based for SQUIRLS) would predict negative variants as splicing-altering. Indeed, we observed a large proportion of false positives for these tools in the pseudoexon-activation group when using a single reference threshold for evaluation (0.99 for TraP, 0.95 for regSNP-intron and 0.64 for SQUIRLS, Supplementary Table S7). Conversely, deep learning based methods such as SpliceRover, DSSP and Spliceator may rely too much on the near-splice site features (despite using larger sequence contexts), hence the poor performance observed.

### Donor-downstream variants

This category refers to all splicing-altering intronic variants located downstream of the cryptic splice donor event (N=148). Negative variants (N=169) are located downstream of annotated exons and were shown experimentally to have no impact on splicing outcomes (Table 2, Supplementary Table S6). As before, we included SpliceRover, Spliceator and regSNP-intron in the analysis. DSSP was excluded since it predicts splice sites at fixed positions in the input, but in this category, variant positions with respect to the cryptic splice donor are variable. SpliceAI excelled on the subset of variants triggering pseudoexon activation with an AP score of 0.95, followed by ConSpliceML, Pangolin and CI-SpliceAI, all with performance values above 0.9 (Figure 3B, Supplementary Figure S4H, Supplementary Table S7). Regarding the partial intron retention subset, SpliceAI and Pangolin performed the best (AP=0.87), with a larger difference for the tool ranked third, CI-SpliceAI (AP=0.814, Supplementary Figure S4I). Again, these results demonstrate the superiority of SpliceAI-derived approaches versus standard methods that engineer domain-specific features to score intronic splicing variation.

### All regions combined

Next, we combined all the datasets to inspect the global performance of each major variant group. Eight methods were able to score all types of splicing variants in any intronic region. Those tools were SpliceAI, Pangolin, ConSpliceML, CI-SpliceAI, AbSplice-DNA, SPiP, TraP, and SQUIRLS (Figure 3C). Except for AbSplice-DNA, which scores intronic variants located up to 100bp away from splice junctions used in any GTEx tissue, all the methods were designed to score any given position in introns.

SpliceAI and Pangolin consistently ranked highly for several datasets (Figure 3C), yet SpliceAI is slightly superior given the results obtained by Pangolin for intron retention variants (exonic-like and new splice donor regions). CI-SpliceAI, AbSplice-DNA and ConSpliceML were fair alternatives, especially for variants that create new splice donors. SPiP was particularly inadequate for exonic-like variants, but was the best non-deep learning-based method for the remaining categories (Figure 3C).

Overall, we observed a trend of pseudoexon-activating variants being predicted more accurately than partial intron retention variants (Figure 3C, Supplementary Figure S6A). However, when analyzing each tool individually, this trend did not reach statistical significance (Supplementary Figure S6B).

### Assessing interpretability

We were interested in the extent to which these state-of-the-art tools give additional information to the user, besides the prediction. Among the tools that predict across whole introns, SQUIRLS and SPiP are the only ones designed to provide some interpretation of the outcome. SQUIRLS can generate HTML reports with short descriptions of why the model predicts pathogenicity and displays the contribution of each feature to the outcome. In addition, it draws figures to show the variant effect in the sequence context surrounding the variant. SPiP provides short interpretation tags describing the molecular consequences of the variants along with confidence intervals for the probability that the variant impacts splicing. We devised a procedure to evaluate how accurate the explanations are against the biological ground truth (see Methods). We use the splicing-associated deep intronic pathogenic variants analyzed above (N=243, Figure 2) to measure the quality of the explanations. SPiP and SQUIRLS correctly predicted 179 and 124 variants, respectively, and those were selected for downstream analysis. We found that SQUIRLS and SPiP were able to provide correct explanations (within the limits of each tool) for a considerable fraction of the correctly predicted pathogenic variants, yet for many others no explanation was found (45 variants with SPiP, 19 variants with SQUIRLS, Figure 4A). Looking at the prediction score distribution for each explanation category, we observed that the variants with no explanation have the lowest scores (Supplementary Figure S7). On the other hand, correct explanations are spread across the full score range. Interestingly, SPiP explanations that are not informative (complex events evolving multiple splicing motifs) have the highest median score range, showing that strong deleterious effects are not necessarily easier to explain.

**Figure 4:**
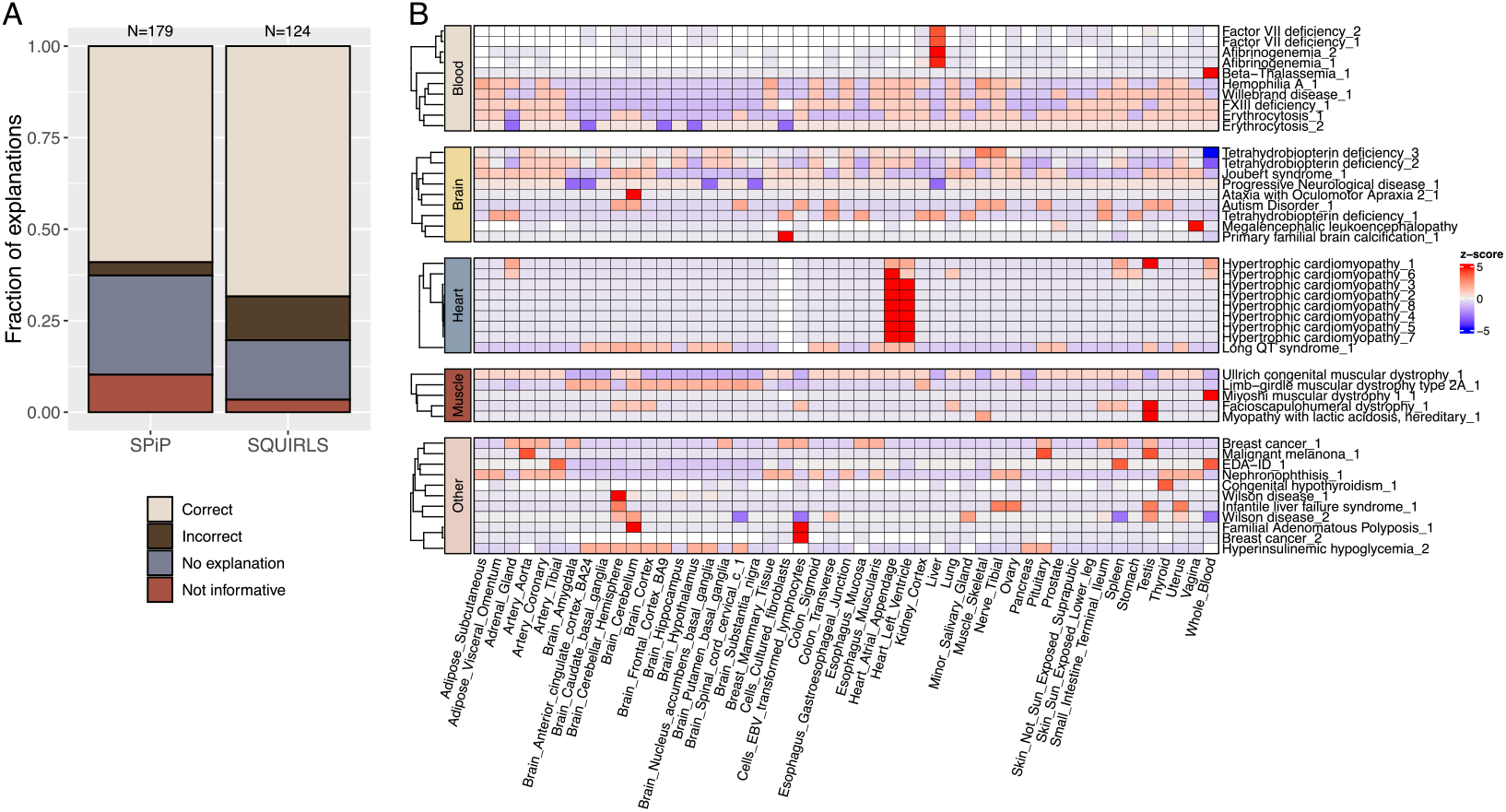
Information provided by the tools beyond the prediction score. **A** - Assessing the quality of the explanations for SPiP and SQUIRLS. Within each bar, the height of each category represents the fraction of variants assigned to the given explanation quality tag. The numbers above the bars indicate how many pathogenic variants were used (pathogenic variants each tool predicted correctly). **B** - Tissue-specific predictions made by AbSplice-DNA for a set of disease-causing variants associated with a single tissue, according to Human Phenotype Ontology (HPO). Phenotype names are displayed in rows, GTEx tissues predicted by AbSplice-DNA are in columns. High z-scores represent tissues for which the variant effect is stronger compared to other tissues.

### Predicting splicing changes across tissues

Of the tools evaluated in this study, Pangolin and AbSplice-DNA can both predict splicing outcomes in a tissue-specific fashion. We decided to use AbSplice-DNA alone for this analysis. Pangolin was trained on sequences and splice site usage levels from four tissues across four species (human, rhesus macaque, mouse and rat). However, the default settings of Pangolin are tissue-agnostic and it requires additional customizations to get tissue-specific variant effect predictions. On the other hand, AbSplice-DNA provides pre-computed tissue-specific predictions. Moreover, it combines tissue-specific splicing annotations created from GTEx data with DNA-based prediction models, enabling it to predict variant effects in more tissues (49). We aimed to evaluate whether AbSplice-DNA predictions of disease-causing variants are enriched for the tissues that are most strongly affected by the disease.

Using the splicing-associated variant dataset described above (N=243, Figure 2), we determined, when possible, the GTEx tissue most closely associated with the given disease based on the HPO [83] (see Methods). We selected the 154 variants that AbSplice-DNA predicted correctly, which excluded all the variants causing two of the most common diseases in our dataset: Becker muscular dystrophy and Duchenne muscular dystrophy (Supplementary Table S9). In addition, 65 variants were not evaluated since they were not assigned to any particular tissue (e.g. systemic diseases, or diseases affecting tissues not represented in GTEx, such as the retina). Considering disease variants associated with only one GTEx tissue, we observed enrichment of the expected tissues to a limited extent (Figure 4B, Supplementary Figure S8A). For example, Hypertrophic Cardiomyopathy variants were highly enriched in the heart tissue, an Ataxia with Oculimotor Apraxia variant was predicted to affect the cerebellum and a Congenital hypothyroidism variant was enriched for thyroid. Interestingly, variants associated with blood disorders (Factor VII deficiency and Afibrinogenemia) have the highest prediction scores in the liver, which is not unexpected, since the liver plays a crucial role in the production of clotting factors, including factor VII and fibrinogen (Figure 4B). However, other tissue-specific predictions had unclear interpretations, such as the enrichment of testis in muscular dystrophies, the brain cerebellum in Adenomatous Polyposis (associated with colon and rectum, Figure 4B), or the skeletal muscle in Fabry disease (primarily linked to other tissues such as heart and kidneys, Supplementary Figure S8A). In addition, 38 variants displayed the same score across all tissues, which does not reflect the expected biology, especially for some diseases associated with a single tissue (Supplementary Figure S8B).

## Discussion

We have performed a comprehensive benchmark study of intronic variant prediction, focusing on disease-causing deep intronic variants affecting splicing via pseudoexon inclusion or partial intron retention. Furthermore, we collected and examined variant sets based on their location relative to the splice sites affected by the altered splicing. Finally, we extended our previously developed package VETA [84] to perform benchmarks on intronic regions and have developed a simple utility to prepare input from VCF files for several web-based splicing-related methods.

We used two different datasets to study intronic variants causing human disease. ClinVar is a database that has been widely used for this purpose. Nevertheless, to the best of our knowledge, it has not been used to evaluate performance as a function of distance to the splice sites. Averaging performance across all bins, we found that splicing-associated tools performed the best overall on ClinVar data. Importantly, we observed a decrease in performance immediately after the two splice site positions, with a particularly noticeable decline at a distance of 11 base pairs from the closest splice site. These results demonstrate the extent to which these methods are biased to predict splice site variants, whereas smaller effect-size variants deeper inside the intron go mostly unnoticed. For many of the tools, such as S-CAP or MLCsplice, this is not unexpected, as they were not designed to predict variants in deep intronic regions. In addition, we observed that some of the variants that appear deep-intronic in the canonical isoform are exonic or located close to the splice sites in other transcripts of the associated gene. Therefore, and according to the American College of Medical Genetics and Genomics and the Association for Molecular Pathology (ACMG-AMP) guidelines [85], we recommend considering multiple isoforms when interpreting deep intronic variants, especially when the canonical isoform is not highly expressed in the tissue of interest [86].

Additionally, we curated a diverse set of pathogenic deep intronic mutations that exclusively affect splicing. Tools that predict across all intronic regions, notably SpliceAI-derived models, showed satisfactory performance. Many variants in this dataset generate new splice sites deep within introns, activating pseudoexons. We speculate that sequence-based models that predict splice sites are particularly well suited to predicting this class of variants, likely because the pseudoexons resemble the sequence context of authentic exons [80] that were presented during their training.

To better understand performance differences between classes of variants, we collected a diverse set of experimentally tested splicing-associated variants, and evaluated the tools’ ability to distinguish them from similar non-splice-altering variants. Region-specific analysis revealed substantial differences in performance. We found that variants affecting the BP and exonic-like splicing regulatory elements were among the hardest to predict. Not even the inclusion of tools specifically built for these regions improves results. BPs are known to be highly degenerate, with only the A and the T located two base pairs upstream being highly conserved within the consensus branchpoint motif [87]. In addition, BPs can occur at different positions within the intron, some introns have multiple BP, and the molecular consequences of interfering with the BP can be very diverse [88, 89, 90]. Similarly, predicting variant effects on splicing regulatory elements is challenging. The binding motifs of many splicing factors are highly degenerate or even unknown, and their impact on splicing largely depends on the cell type [91, 92]. Such complexity appears to be better captured by SpliceAI-like deep learning methods than by tools with built-in domain knowledge.

The recent progress achieved through deep learning models that work as black boxes has raised concerns about their deployment in sensitive domains such as healthcare [93]. Because practitioners are interested in understanding how these AI systems make decisions, we assessed the capacity of these models to provide interpretable outputs when predicting disease-causing variation associated with splicing defects. Although most sequence-based models, such as SpliceAI, provide some information beyond the prediction score, namely the distance of the variant to the affected mRNA position, it is only possible to obtain insight into the inner workings of the model by applying external explainability techniques [94]. On the other hand, SQUIRLS and SPiP are intrinsically more interpretable by design. The models were frequently able to correctly identify the type of splicing alteration, yet they still fail to propose higher-order mechanistic hypotheses for such predictions. In addition, these models suffer from an accuracy-interpretability trade-off since the performance across evaluations was lower than that of black box models.

Another promising research avenue is the prediction of splicing abnormalities in a tissue of interest, which AbSplice-DNA offers. The model could accurately detect some tissue-specific differences relevant to human disease, yet it was unreliable for the majority of variants. Nonetheless, we acknowledge that the introduction of SpliceMaps [95], which provide information on splice site usage across GTEx tissues, combined with RNA-Sequencing of clinically accessible tissues (CATs), is expected to enhance the prediction of functional intronic variants [25], particularly in diseases where the splicing landscape of the relevant non-accessible tissue is appropriately represented by one of the CATs [96].

### Practical recommendations

We advocate using deep learning-based solutions if the main goal is to predict as accurately as possible. SpliceAI and Pangolin consistently ranked high for intronic variants associated with splicing, both for the prediction of pathogenicity and of altered splicing. If usability is a concern and users do not have a large number of predictions to make, SpliceAI is preferred since the Broad Institute has made available a web app for the task (https://spliceailookup.broadinstitute.org/). If the number of variants makes it unfeasible to use the SpliceAI-lookup app, CI-SpliceAI is a good alternative since its web app allows the input of multiple variants in a VCF-like format (https://ci-spliceai.com/). Practitioners, however, may suspect that splicing is not the mechanism disrupted by a particular mutation. In this scenario, we recommend prioritizing intronic variants using CAPICE since it was the whole genome predictor with the best results with ClinVar data, although its performance was limited.

Region-specific splicing benchmarks revealed additional insights for tool usage. We recommend using Pangolin to prioritize variants in branchpoint regions (−18 to -44 bp upstream of splice acceptor). LabRanchoR was the best branchpoint-specific model in our evaluation and can also be considered. SpliceAI and Pangolin were the most effective at scoring acceptor-associated variants (splice acceptor-creating or polypyrimidine tract variants upstream of cryptic splice acceptors). Including other sequence-based deep learning models that use smaller sequence contexts did not provide additional value. Intronic variants affecting splicing regulatory elements within cryptic exons are hard to predict. We endorse using SpliceAI with larger windows surrounding the variant site (setting the distance parameter to the maximum). In addition, classical approaches such as HEXplorer might come in handy for specific cases, such as assessing the potential impact of a variant on exon-defining regulatory motifs. Finally, SpliceAI-inspired models (Pangolin, CI-SpliceAI) and models that incorporate SpliceAI predictions as features (ConSpliceML, AbSplice-DNA) can effectively predict new splice donor and donor-downstream variants. However, to keep the number of different tools to use to a minimum, we suggest using the original SpliceAI model.

Nonetheless, it is noteworthy to mention the impacts of using pre-computed scores as the strategy for variant prioritization. The current version of SpliceAI pre-computed scores (v1.3) does not include predictions for insertions and deletions larger than 1 and 4 nucleotides, respectively. In addition, the limit of 50 bp as the distance around the variant site to extract variant effects prevents SpliceAI from identifying other variant classes, such as exon skipping, when the variant exerts its effects at more than 50 base pairs from the affected exon.

Finally, when interpretable outcomes are important, we recommend SQUIRLS as the method of choice. This software is well-documented and generates HTML reports that practitioners can intuitively inspect. Nonetheless, it is important to note that SQUIRLS should not be solely relied upon, as it is not as performant as other models.

### Final remarks

We comprehensively assessed functional intronic variation occurring far from annotated splice sites. As a result, we make available to the community region-specific variant sets that can be used to evaluate new models on variants whose molecular consequence is known. This will assist developers in identifying potential limitations of the model and highlighting variant types that it is more prone to fail on. Additionally, we encourage developers to make their models publicly available by sharing them on open-source platforms to facilitate their reuse [42, 97].

Sequence-based models based on Convolution Neural Networks architectures are still the state-of-the-art approach for splicing variant prediction. However, the artificial intelligence field is rapidly evolving, and we have seen the emergence of Transformer-based architectures being applied for other variant effect prediction tasks, e.g., effects on gene expression [98] or on protein function using large protein language models [99]. As a result, increasingly complex models are expected to effectively tackle open questions in splicing regulation, such as better capturing the synergistic effects of splicing regulatory elements. However, the community must be aware of the possible implications these models bring, such as a lack of transparency and decreased ability to generate mechanistic hypotheses.

## Methods

### Data collection and variant annotation

We employed the same variant annotation procedure for all the variants collected for this manuscript (datasets described below). We used Ensembl VEP v107 ([100]) for the task and transcript annotations were added accordingly (with ‘–per_gene –pick_order ccds,canonical,biotype,rank –no_intergenic –gencode_basic’ set). We used variants in the GRCh37 genome build simply because several of the tools we include in the manuscript do not support the GRCh38 genome build. Nonetheless, we provide all the datasets and predictions in both GRCh37 and GRCh38 (via liftOver) versions.

### ClinVar

We downloaded ClinVar v202204 and selected all the SNVss for downstream analysis. We kept variants with Pathogenic and Benign assignments (‘CLNSIG’== Pathogenic or Likely_pathogenic or Benign or Likely_benign). We identified intronic variants based on Ensembl VEP annotations: only variants with at least one intronic consequence (‘INTRON’== 1) in a protein-coding transcript (‘BIOTYPE’= protein_coding) were retained. Additionally, we excluded variants with exonic annotations in any other gene (‘EXON’≠ 1). To avoid being overly conservative, we added variants that Ensembl VEP annotated as being outside the gene body for the picked consequence (‘Consequence’== TF_binding_site_variant or downstream_gene_variant or up-stream_gene_variant or regulatory_region_variant), but that are annotated with intronic ontology terms in the ‘MC’field in the original VCF. To minimize labeling errors, we excluded variants with less than one confidence star. Finally, to ensure that the number of benign variants did not exceed 50,000 (and therefore avoid the dataset being excessively unbalanced), we selected all higher-confidence benign variants (with two or more stars, N=13,093) along with 36,907 randomly chosen one-star variants. The final ClinVar intronic dataset size amounted to 18,608 pathogenic and 50,000 benign variants.

### Disease-causing intronic variants affecting RNA splicing

This dataset refers to a high-quality variant set that we carefully curated to comply with the following criteria:

- Variant must locate at more than 10bp from the nearest splice site.
- Variant was experimentally proven to affect normal RNA splicing.
- Variant does not necessarily lead to pseudoexon activation.

The previous curation effort from our lab ([19]) was updated for this manuscript to include a comprehensive set of intronic variants identified after 2017. Therefore, the positive (disease-causing) set of variants used in this benchmark totals 243 (81 from Vaz Drago et al. (2017) and 162 from the new curation effort).

We used gnomAD v2.1 to generate a matched control set. First, we extracted all gnomAD variants occurring in a window of 500bp surrounding the variants in the positive set and selected common records with a frequency higher 0.01 (1%) in the population, resulting in 1128 variants. Then, we ran Ensembl VEP as previously described and retained the intronic variants annotated as occurring in one of the 148 unique genes of the positive set (N=1091). Moreover, we kept variants absent in ClinVar having the VCF filter field as ‘PASS’(N=546). Finally, we randomly sampled 243 from this set.

### Variants that affect RNA splicing

The third main dataset refers to variants that affect different mechanisms of splicing regulation, which may or may not lead to disease. We defined different molecular categories based on the location of the variant relative to the abnormal splicing event. We focused on deep intronic variants that lead to partial intron retention or pseudoexon activation. Exceptionally, we included variants that affect the branchpoint signal (thus, closer to annotated splice acceptors) that include other types of splicing alterations such as exon skipping. We defined each category as follows:

- *Branchpoint-associated*, for those variants occurring between -18 and 44bp upstream of an annotated or cryptic splicing acceptor site, as used in [31], that may affect the branchpoint signal.
- *Acceptor-upstream*, referring to any variant that locates between -2 and -18bp upstream of the cryptic splice acceptor (including the polypyrimidine tract).
- *New splice acceptor*, denoting the variants that occur at the cryptic splice acceptor positions, including the first nucleotide of the cryptic exon.
- *Exonic-like*, for any variant occurring within the cryptic exon (pseudoexon or partially retained intron).
- *New splice donor*, composed of variants located at the cryptic splice donor positions, including the last position of the cryptic exon.
- *Donor-downstream*, referring to any deep intronic variant that locates at a distance of more than 2bp from the activated cryptic splice donor.

We used data produced or gathered from multiple studies to assign variants to each category (Table 2). While splicing-altering variants were straightforward to assign (based on source data, the functional consequence, and distances to the splicing element considered), we distributed the non-altering variants such that they resembled as best as possible the spatial distribution of the positive sets. Hence, we assigned the negative variants taking into account two levels of information: the primary group (partial intron retention, pseudoexon activation) and the region category.

To keep in line with the expected biology, we assigned the variants that were within a defined distance to a splice site to the partial intron retention group and deeper intronic variants to the pseudoexon group. We used different distance thresholds for splice acceptors and donors (150bp and 20bp, respectively) so that the datasets were reasonably balanced. As for the region category, we defined negative variants occurring between 18 and 44bp upstream of an annotated splicing acceptor as branchpoint-associated variants. We assigned as acceptor-upstream or donor-downstream the remaining intronic variants according to whether they were located upstream or downstream of the nearest annotated splice site. Because pseudoexons tend to resemble authentic exons [80], we exceptionally assigned exonic variants that did not change inclusion levels of tested exons [81] as controls for the Exonic-like category.

Lastly, we generated control datasets for the new splice site categories. Splice site variants (located at one of the dinucleotide positions) that were experimentally tested to not affect splicing are not easily accessible. Therefore, to mimic the positive set, we looked for common deep intronic SNVss (> 5% in gnomAD v2.1) in protein-coding transcripts that generate the most common 5-mer acceptor motif CAGGT in the human genome [14] through a mutation in the core splice site dinucleotide. We randomly selected 67 variants to match the number of positive new splice acceptor variants exactly. We employed the same procedure for the new splice donor variants, where we kept the variants that generate the most common 6-mer donor motif GGTAAG in the human genome [14] at the GT position. Finally, we selected 171 variants at random to match the number of positive new splice donor variants. We confirmed using Snaptron [101] that in GTEx data, there is no evidence that a splice junction is used at the variant intervals, except for two variants.

### Prediction tools

We selected an extensive list of prediction tools for evaluation. The single criterion for the inclusion of a tool was that it had to be designed to predict (at least partially) intronic variation. When available, we used precomputed scores to annotate our variant sets (from dbNSFP v4.0b1 [102], UCSC genome browser [103], Zenodo or tool website). Otherwise, we ran the models directly following the developer’s instructions. For a subset of splicing-related tools (MMSplice, HAL, kipoiSplice4, MaxEntScan), we employed kipoi v0.8.6 [42] to get predictions. We additionally included splicing-related tools that predict specific splicing signals (e.g., BP) and are not necessarily targeted to predict pathogenicity. Because most of these tools do not score variants by design, and some require using a web-based portal, we developed a simple utility to prepare their input given a VCF file. Moreover, we created a script for each tool to process the raw output into a final prediction score to be included in a VCF file. The package is available at https://github.com/PedroBarbosa/Prepare_SplicingPredictors. We annotated the final VCF files with all the predictions using vcfanno v0.3.2 [104]. We describe all the tools, their reference thresholds and how we ran them in Table 1.

### Performance evaluation

We used VETA v0.7.4 [84] to perform all the performance evaluations. We extended VETA’s feature set by implementing a new mode (“–do_intronic_analysis”) targeted to intronic variants. This option assigns intronic variants into distance bins based on their distance to the closest splice site. Accordingly, VETA seamlessly integrates per-bin analyses, allowing automatic inspection of how tool performance varies as one moves deeper into the intronic space. Moreover, VETA includes an *interrogate* mode that ranks candidate variants according to tool predictions, facilitating the downstream variant interpretation task in whole genome and exome studies.

In this manuscript, we employed different metrics according to the nature of the dataset and the goal of the analysis. Despite this, VETA generated confusion matrices for all tools. True Positives (TP) indicates the number of pathogenic/splicing-altering variants that a tool correctly predicts as pathogenic (or splicing-altering). True Negatives (TN) is the number of benign (or non-splicing-altering) variants that a tool scores as such. False Negatives (FN) refers to the number of true pathogenic/splicing-altering variants that a tool predicts to be benign/non-splicing-altering. Finally, False Positives (FP) stands for the number of benign/non-splicing-altering variants that a tool scores as pathogenic (or splicing-altering). For ClinVar data, we ranked variants based on the F1-score, given the unbalanced nature of the data (much more intronic benign variants than pathogenic). Because some tools do not score deep in the introns (missing data), we weighted the F1-score with the prediction coverage: 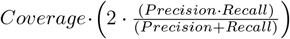 where 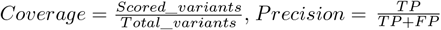 and 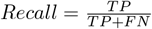 For balanced datasets, we ranked tools using a slight variation of the Matthews Correlation Coefficient (MCC) 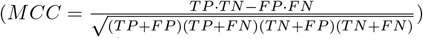 that normalizes the metric range between 0 and 1 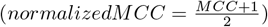 We weighted the normalized MCC values with the prediction coverage (*weighted*_*normalized*_*MCC* = *Coverage normalizedMCC*). Additionally, we employed ROC and PR Curves for the comparisons that measure performance at multiple threshold values. To summarize such analyses, we used the auROC and AP metrics, respectively.

### Further inspection of deep intronic variants in ClinVar

We selected ClinVar variants assigned to the “501-1000” and “+1000” intronic bins and used VEP to perform reannotation. We ran VEP without picking any consequence (“–per_gene” and “–pick_order” were not set), meaning that all transcript consequences associated with each variant were retained. We employed a filter to only keep annotations of protein-coding transcripts. Then, we assigned each variant to one of four categories, according to the overlap configuration of transcripts belonging to the gene associated with the variant: if a variant is exonic in another overlapping transcript, we termed it as “Exonic”; if a variant is located at a shorter distance from the splice site in any other transcript, we assigned the category “> 1 transcript (smaller offset)”; if the distance to the closest splice site remains the same for all transcripts overlapping the variant, we assigned the variant to the “> 1 transcript (smaller offset)” category; lastly, if no other transcript overlapped with the variant (besides the one used in the analysis), we set it to the “No other transcript” category.

### Threshold analysis for deep intronic variants

To derive clinically applicable prediction thresholds for deep intronic variants, we employed the same strategy we recently described [84]. Briefly, for each tool, we applied the F-Beta formula (at three different Beta values) over 100 threshold values uniformly distributed between the range of scores. The threshold that maximized the F-Beta function was selected. To evaluate the reliability of the adjusted thresholds, we used a bootstrapping procedure, where we kept the same ratio of pathogenic and benign variants as in the original dataset in each bootstrap. This analysis was conducted using VETA, with the options “–do_threshold_analysis” and “–bootstrapping” enabled.

### Combining tools to build a Random Forest model

We combined tool predictions to create standard scikit-learn [105] classifiers to distinguish splicing-altering from non-splicing-altering variants. Tools with more than 30% missing data in any of the class labels were discarded. Then, missing values were imputed using the median value along each column. We split the variants into a training set (80%) and a test set (20%) in a stratified manner. We performed a grid search with cross-validation to find the best set of parameters for the classifier, and used the best model to evaluate performance on the test data. We implemented this analysis in VETA with the option “–do_machine_learning”.

### Assessing quality of explanations for SPiP and SQURLS

For this task, we employed the dataset of pathogenic splicing variants used throughout the study. It includes variants from our curation plus variants from Vaz-Drago et al. [19] because the molecular mechanism for the splicing defect is known for almost all records (Supplementary Table S3). For each tool, we ran VETA in the *interrogate* mode (with “*–labels Pathogenic*” set) to list the variants correctly predicted by each tool using the threshold calibrated for non-canonical intronic variation (SPiP > 0.009 and SQUIRLS > 0.016). We removed variants for which the ground truth information was not available (e.g.,pseudoexon-activating variants that lack details of the location of the variant concerning the cryptic event).

For SPiP, we parsed the output so that the interpretation tag, confidence interval and original score were retrieved (3rd, 4th and 5th fields after splitting predictions by “|”). We assigned variants with an “NTR” tag (low probability of affecting splicing, yet correctly predicted as pathogenic according to the calibrated threshold) to the “No explanation” category. Variants affected by multiple splicing motifs (according to SPiP, the “Alter by complex event” tag) were given the “Not informative” explanation category. Then, for each of the remaining SPiP interpretation tags we classified the explanation as correct if they matched the ground truth information:

- “Alter BP” for variants associated with the branchpoint signal, else incorrect.
- “Alter by create new Exon” for variants that trigger pseudoexon activation, else incorrect.
- “Alter by create New splice site” for variants that create a new splice site or activate a nearby existing cryptic splice site, regardless of the variant leading to pseudoexon activation or partial intron retention, else incorrect.
- “Alter ESR” for intronic variants occurring within the boundaries of a new pseudoexon, else incorrect.
- “Alter by MES (Poly TC)” for polypyrimidine tract variants, else incorrect.

As for SQUIRLS, we ran the model for the subset of pathogenic variants correctly predicted by the tool using “–output-format html” and “–n-variants-to-report 125”. Afterwards, we manually inspected the HTML report generated to derive structured interpretations for each variant: “Not informative” if the short description of the variant effect was not generated; “No explanation” if SQUIRLS did not produce any description or figure for the variant; “New cryptic acceptor” and “New cryptic donor” if SQUIRLS described the creation of a new splice site and the variant was located at one of the splice site positions (based on the Sequence trekker figure) defined in this manuscript; “Activate cryptic acceptor” and “Activate cryptic donor” if SQUIRLS described the creation of a cryptic splice site and the variant was located outside of the splice site positions (based on the Sequence trekker figure). Because SQUIRLS does not predict the exact molecular effect of a splicing variant, we ignored the predicted number of bases affecting the coding sequence as this was not applicable for pseudoexon-activating variants. After manually inspecting the HTML report and generating structured explanations, we classified the explanation as correct if it matched the ground truth information:

- “New splice acceptor” for variants that create a new splice donor, else incorrect.
- “New splice donor” for variants that create a new splice acceptor, else incorrect.
- “Activate cryptic acceptor” for variants located upstream of an existing cryptic splice acceptor and not associated with the branchpoint signal, else incorrect.
- “Activate cryptic donor” for variants located downstream of an existing cryptic splice donor, else incorrect.

### Tissue-specific predictions by AbSplice-DNA

We selected the maximum AbSplice-DNA prediction for any tissue to evaluate model performance. Here, to measure how accurate the tissue-specific predictions of the model are, we used the same dataset as for the interpretability section. We ran VETA in the *interrogate* mode (with “’*–labels Pathogenic*” set) to list the variants correctly predicted by AbSplice-DNA using the threshold adjusted for non-canonical intronic variation (>0.004). Then, for each variant, we gathered information about the tissues associated with the disease by searching the HPO [83] with the given OMIM disease identifier. We strived to assign tissue names that matched the GTEx tissues used by AbSplice-DNA. Disease-causing variants affecting tissues not represented in GTEx (e.g. retina) were discarded. Additionally, variants causing systemic diseases (e.g. Marfan syndrome), or diseases returning ambiguous HPO terms were excluded.

## Code and data availability

All datasets and code to reproduce the results are available at https://github.com/PedroBarbosa/paper_intronic_benchmark. The benchmarks performed rely on VETA, an open-source tool to compare prediction tools from VCF files (https://github.com/PedroBarbosa/VETA). We also developed a simple utility to generate correct inputs for several sequence-based splicing-related methods from VCF files, available at https://github.com/PedroBarbosa/Prepare_SplicingPredictors.

## Author’s Contributions

Conceptualization: P.B., R.S., M.C-F., A.F; Funding acquisition: P.B., M.C-F, A.F.; Data curation: P.B.; Investigation: P.B.; Methodology: P.B; Resources: P.B., M.C-F; Software: P.B, A.F.; Supervision: R.S., M.C-F., A.F.; Visualization: P.B., Writing - original draft: P.B.; Writing - review & editing: P.B., R.S., M.C-F., A.F.;

## Supplementary figures

**Supplementary Figure S1:**
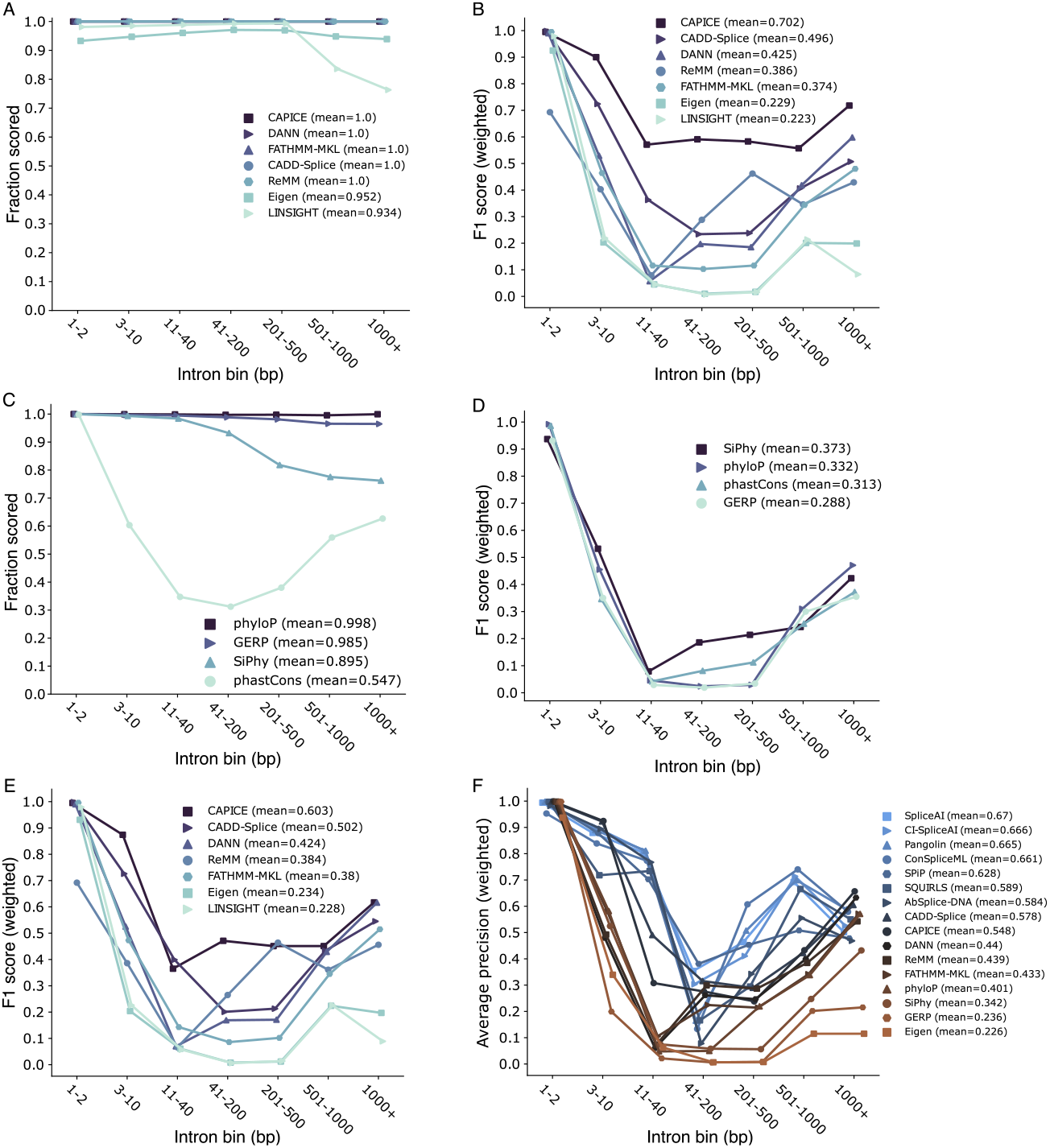
Intronic variant prediction in ClinVar for conservation scores and whole genome predictors at each intronic bin. **A** - Fraction of variants scored for whole genome predictors. **B** - Performance of whole genome predictors. **C** - Fraction of variants scored for conservation scores. **D** - Performance of conservation scores. **E** - Performance of whole genome predictors after removing variants used to train CAPICE (54,117 variants analyzed). **F** - Performance for all tools with sufficient predictions to perform PR Curves in all intronic bins. Evaluations were performed using the dataset without the variants used to train CAPICE (N=54,117).

**Supplementary Figure S2:**
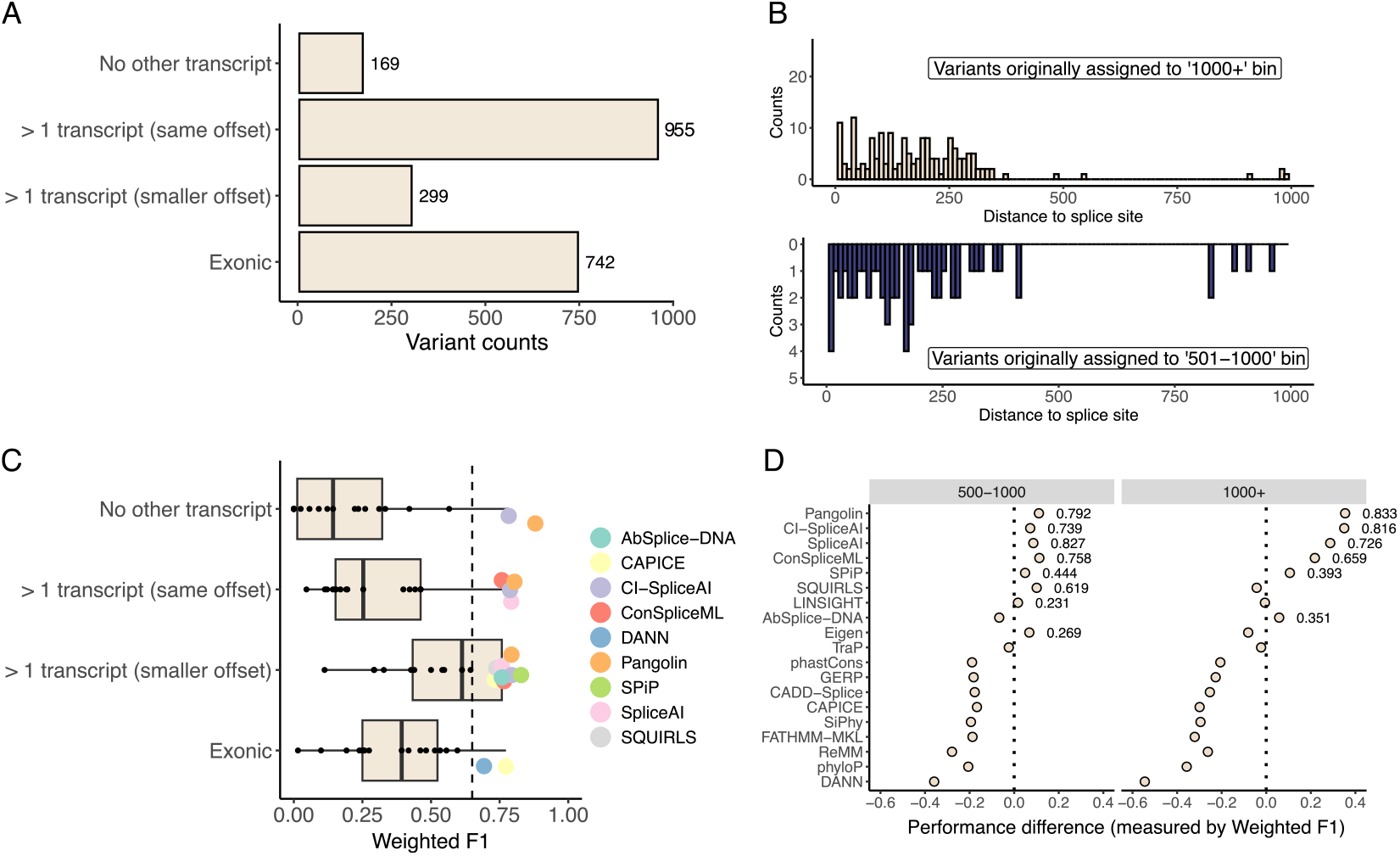
Inspection of ClinVar intronic variants originally assigned to the “501-1000” and “1000+” intronic bins. **A** - The number of variants assigned to each category. The term “No other transcript” refers to all variants that do not have any other transcript of the same gene overlapping with them, besides the transcript originally used (N pathogenic=13, N benign=156). “> 1 transcript (same offset)” refers to variants that overlap with more than one transcript of the same gene but do not have any other transcript where the variant is closer to the splice site than in the original transcript used in the analysis (N pathogenic=57, N benign=898). “> 1 transcript (smaller offset)” refers to variants that overlap with more than one transcript of the same gene, and have at least one other transcript in which the variant is closer to the splice site than in the original transcript used in the analysis (N pathogenic=32, N benign=267). “Exonic” refers to variants that overlap with more than one transcript of the same gene, and have at least one other transcript where the variant is exonic (N pathogenic=150, N benign=592). **B** - Distribution of the updated intronic distances to the closest splice site for variants assigned to the “> 1 transcript (smaller offset)” category. **C** - Tool performance (measured with weighted F1 score) for each individual category. Splicing tools that are not designed to predict these regions were excluded (MaxEntScan, HAL, SPIDEX, dbscSNV, S-CAP, kipoiSplice4, MLCsplice, MMSplice). Tools with performance higher than 0.65 are highlighted. **D** - Differences in performance per deep intronic bin (“501-1000” and “1000+”) after removing variants assigned to the “Exonic” and “> 1 transcript (smaller offset)” categories. Points refer to the weighted F1 difference between this new analysis minus the values obtained with all original variants (displayed in Figure 1C and Supplementary Figure 1B, D). Annotations next to points refer to the weighted F1 scores in the new analysis for the tools whose performance difference is positive.

**Supplementary Figure S3:**
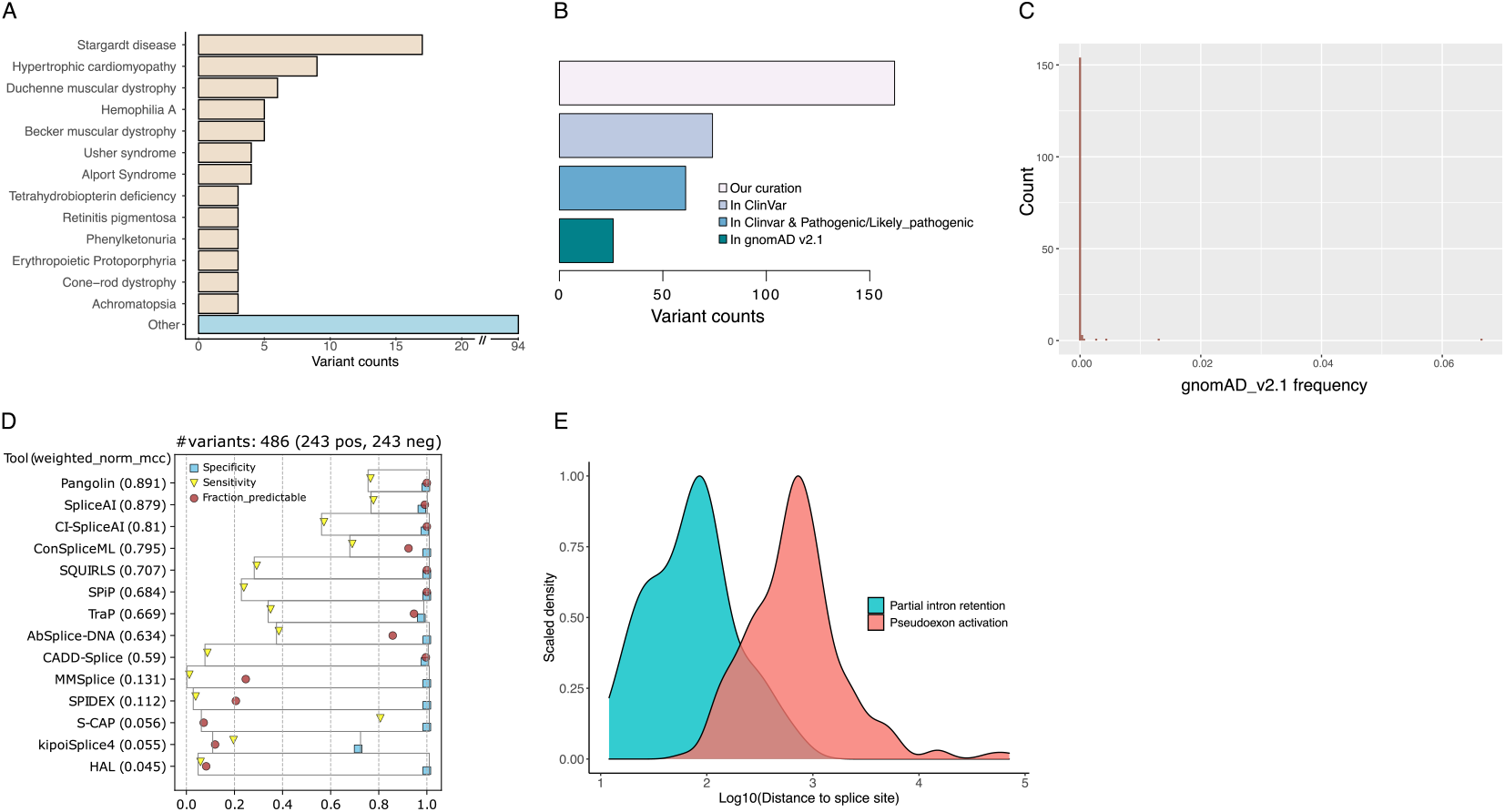
Manually curated dataset of pathogenic intronic variants disrupting RNA splicing. **A** - Number of variants collected per phenotype. Diseases with less than 3 variants were assigned to the ’Other’ category. **B**- Number of variants occurring in ClinVar and gnomAD v2.1. **C** - Allele frequency of variants in gnomAD v2.1. **D** - Tool performance using reference thresholds from Table 1. **E** - Distance (Log10) of the variant to the closest splice junction.

**Supplementary Figure S4:**
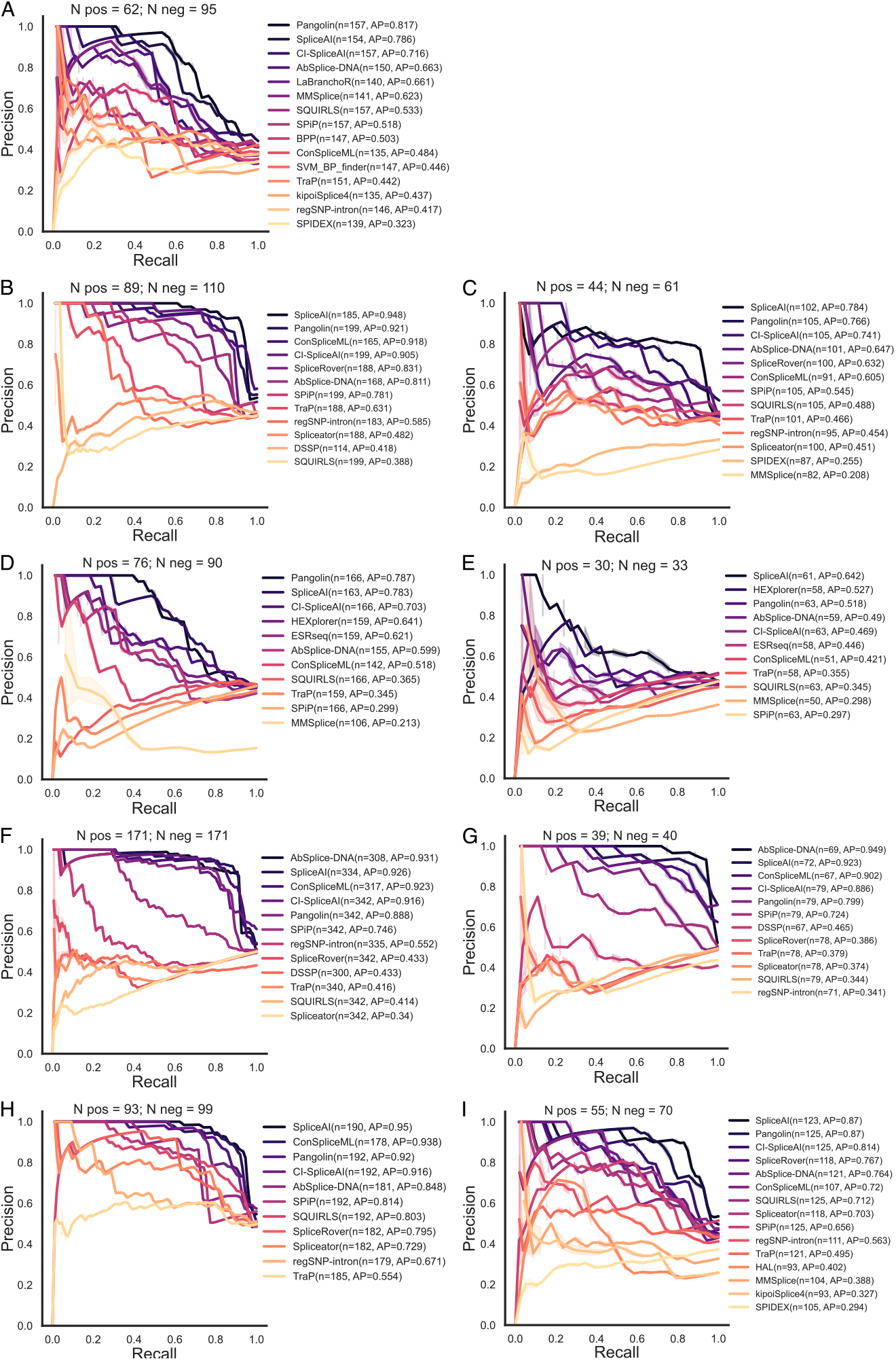
Precision-recall curves for all splicing-altering variants analyzed in a region-specific manner. Tools are ranked by the Average Precision score and the number of predictions made by each tool is displayed in “n=”. The number of variants in each dataset is presented (“N pos” represents the number of positive splicing altering variants; “N neg” is the number of negative splicing variants). Tools with more than 66% of missing predictions or with less than 15 variants in the minority class were excluded from these analyses. **A** - Branchpoint associated variants. **B** - Acceptor-associated variants that trigger pseudoexon inclusion. **C** - Acceptor-associated variants that lead to partial intron retention. **D** - Exonic-like variants that trigger pseudoexon inclusion. **E** - Exonic-like variants that lead to partial intron retention. **F** - Variants that create new splice donors and activate pseudoexons. **G** - Variants that create new splice donors and lead to partial intron retention. **H** - Variants that activate existing upstream cryptic splice donors and trigger pseudoexon activation. **I** - Variants that activate existing upstream cryptic splice donors and lead to partial intron retention.

**Supplementary Figure S5:**
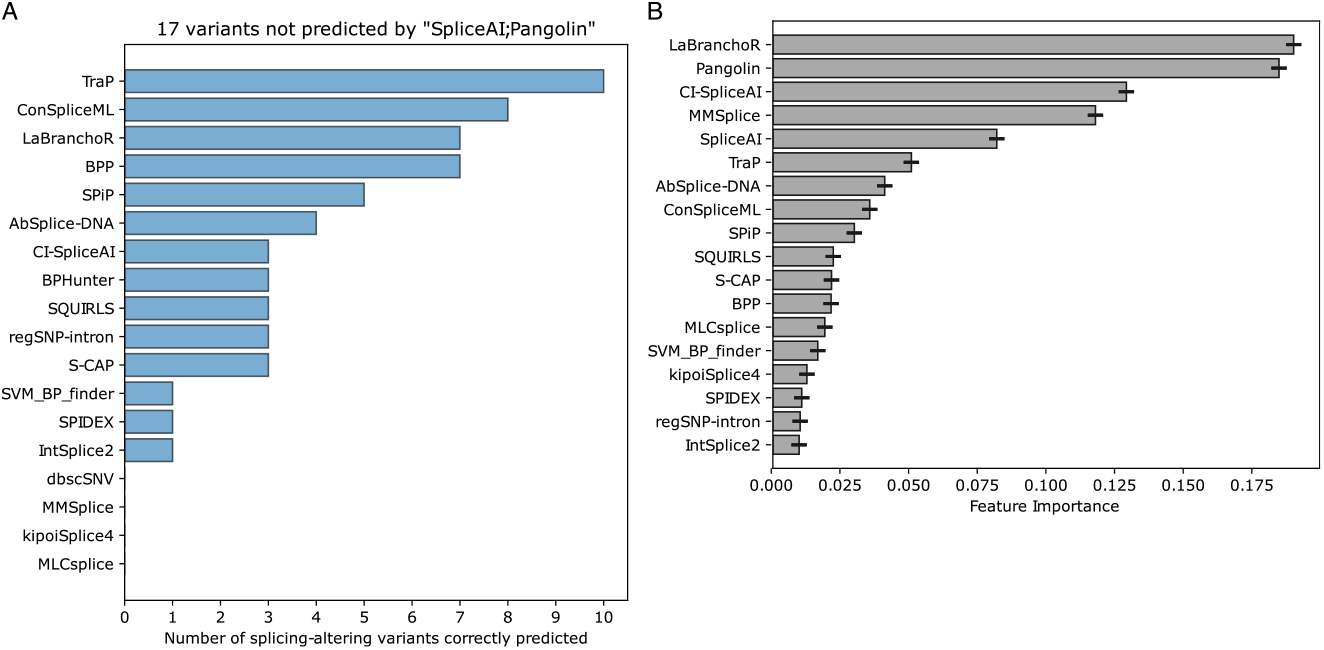
Analysis of branchpoint associated variants. **A** - Number of splicing-altering variants missed by Pangolin and SpliceAI correctly predicted by the remaining tools. **B** - Feature importance (with standard deviation) values for a Random Forest forest model trained on all the branchpoint-associated variants. Importance values represent the average contribution of a feature across all forest trees to reduce the impurity that results from splitting a node in the decision tree.

**Supplementary Figure S6:**
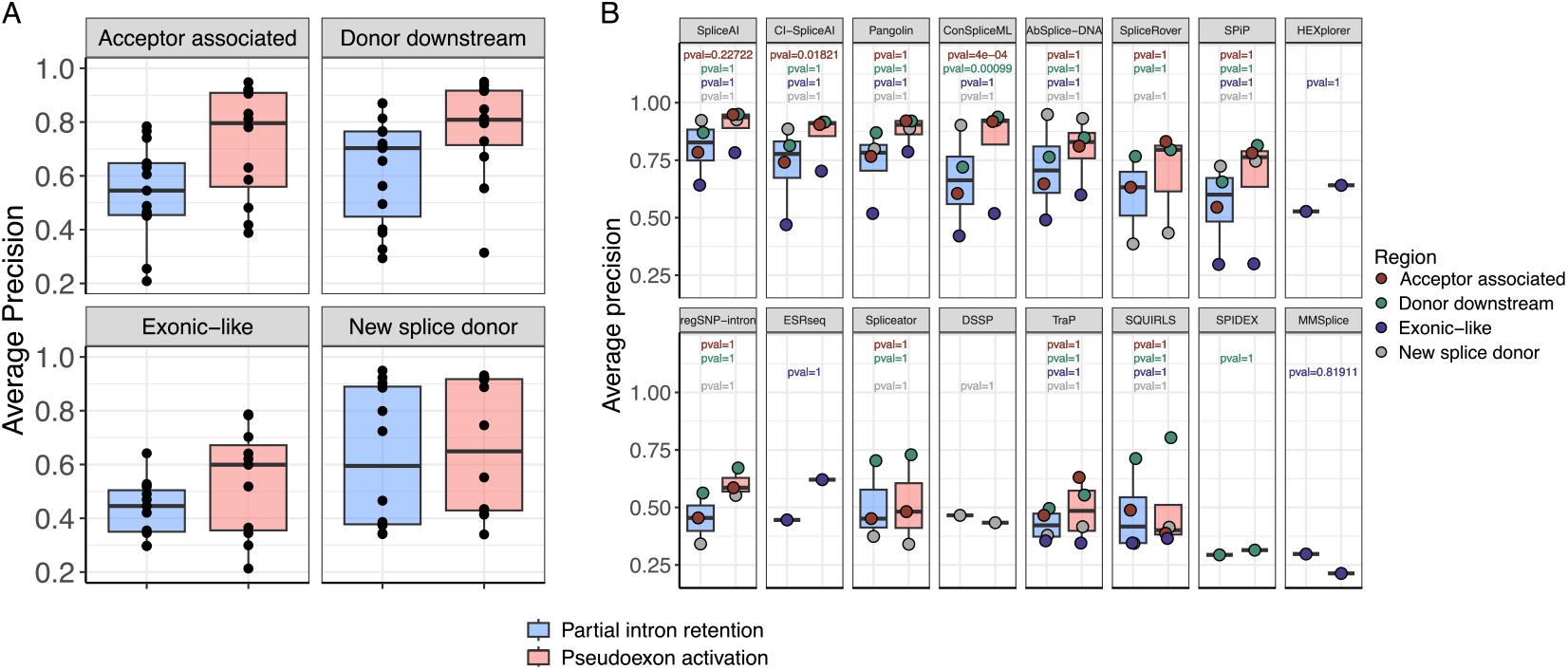
Comparing performance between all pseudoexon activation versus partial intron retention variants collected in this study. **A** - Distribution of the average precisions of the tools for each variant region. **B** - Per-tool average precision distribution across the four variant regions considered. A Fisher’s exact test was conducted separately for each tool and variant region to determine statistical significance for the performance differences between the pseudoexon activation and partial intron retention groups. The true positives plus true negatives were considered successful outcomes, while false positives plus false negatives were considered failures. The p-values displayed in the figure were corrected for multiple comparisons using the Holm method. For each tool, we excluded the variant regions that did not have performance measurements in both groups.

**Supplementary Figure S7:**
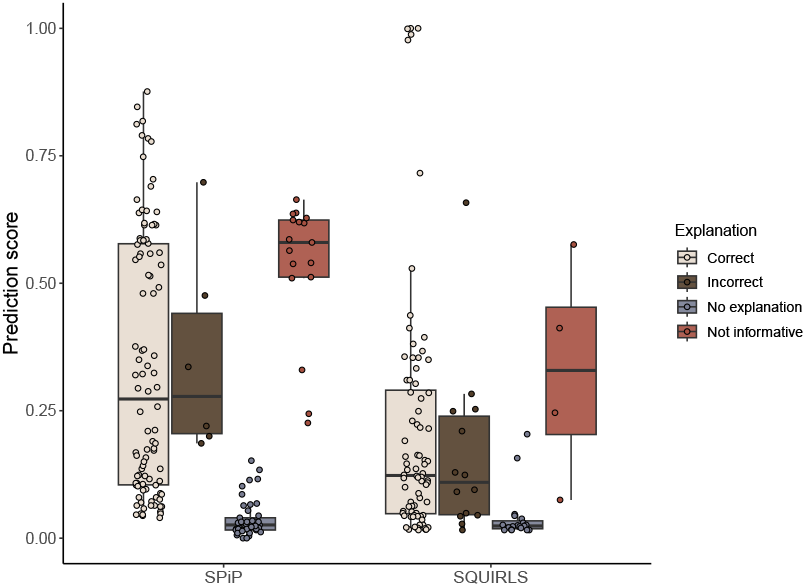
Distribution of SPiP and SQUIRLS prediction values for each of the explanation categories.

**Supplementary Figure S8:**
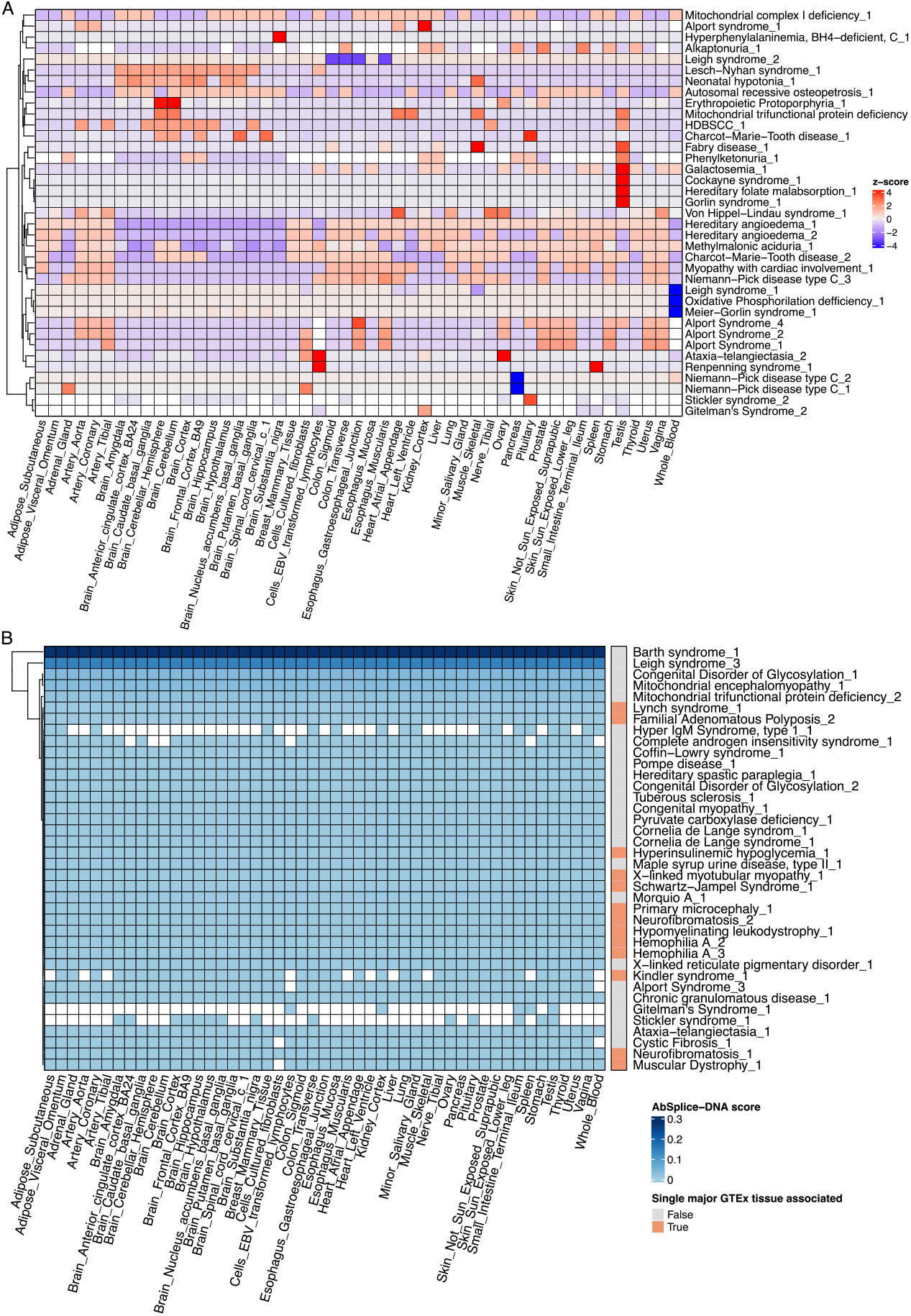
Tissue-specific predictions made by AbSplice-DNA for a set of disease-causing variants associated with aberrant splicing. **A** - Disease variants associated with multiple GTEx tissues that displayed variable scores across tissues. **B** - Disease variants with no tissue-specificity. All tissues got the same AbSplice-DNA score. Disease variants associated with one or more GTEx tissues are displayed in a single heatmap annotation.

